# Structural prediction of chimeric immunogens to elicit targeted antibodies against betacoronaviruses

**DOI:** 10.1101/2023.01.31.526494

**Authors:** Jamel Simpson, Peter M. Kasson

**Affiliations:** Program in Biophysics and Department of Biomedical Engineering, Box 800886, Charlottesville VA 22908; Departments of Chemistry & Biochemistry and Biomedical Engineering, Georgia Institute of Technology, Atlanta, GA 30332; Department of Cell and Molecular Biology, Uppsala University, Box 256, Uppsala, Sweden

## Abstract

Betacoronaviruses pose an ongoing pandemic threat. Antigenic evolution of the SARS-CoV-2 virus has shown that much of the spontaneous antibody response is narrowly focused rather than broadly neutralizing against even SARS-CoV-2 variants, let alone future threats. One way to overcome this is by focusing the antibody response against better-conserved regions of the viral spike protein. Here, we present a design approach to predict stable chimeras between SARS-CoV-2 and other coronaviruses, creating synthetic spike proteins that display a desired conserved region and vary other regions. We leverage AlphaFold to predict chimeric structures and create a new metric for scoring chimera stability based on AlphaFold outputs. We evaluated 114 candidate spike chimeras using this approach. Top chimeras were further evaluated using molecular dynamics simulation as an intermediate validation technique, showing good stability compared to low-scoring controls. Experimental testing of five predicted-stable and two predicted-unstable chimeras confirmed 5/7 predictions, with one intermediate result. This demonstrates the feasibility of the underlying approach, which can be used to design custom immunogens to focus the immune response against a desired viral glycoprotein epitope.

## Introduction

SARS-CoV-2 has caused a global pandemic that has resulted in at least 776 million diagnosed cases of Covid-19 and 7 million deaths as of September 2024^1^. SARS-CoV-2 uses the receptor binding domain (RBD) of its S protein to bind the ACE-2 receptor on host cells, permitting subsequent fusion^2-4^. The S protein has been a focus for vaccine development because of its importance for infection, its display on the viral surface, and its greater immunogenicity compared to other SARS-CoV-2 proteins^5-8^. The RBD is the immunodominant domain of the S protein, eliciting a large majority of neutralizing antibodies^8-10^. However, it also displays the greatest propensity to mutate^11,12^, and RBD mutations in major variants have reduced the effectiveness of current vaccination programs^11,13-17^.

One strategy for overcoming variation in the RBD is to elicit neutralizing antibodies that target other parts of the S protein^18-21^. Two major domains that constitute other targets for neutralization are the N-terminal domain (NTD) and the stalk (S2 domain). Neutralizing antibodies against each have been recovered from patients, demonstrating the potential for SARS-CoV-2 neutralization via epitopes that may be better conserved across variants^20-22^.

Immunization with “chimeric antigens” is one approach that has successfully elicited antibodies against such less immunogenic regions^23-27^. Chimeric antigens can be constructed by exchanging domains from homologous proteins encoded by different viruses to form a novel protein that maintains the overall approximate structure but varies large portions of the primary sequence. Serial immunization with a set of chimeric antigens that hold one domain constant while varying the others can thus amplify the immune response against the conserved domain.

Work by Krammer and colleagues^28^ has tested serial immunization with chimeric antigens against influenza. In this approach, a series of chimeric influenza hemagglutinin proteins maintained the same stalk region in all the chimeras but replaced the head domain with different group 1 influenza virus head domains^28^. Ferrets vaccinated with this combination of chimeras had higher levels of anti-stalk antibody than those vaccinated with standard influenza vaccines^28^. More importantly, the chimeric immunization strategy elicited a broad neutralization response against different influenza subtypes, assessed both in ferret challenge studies^28^ and in neutralization assays as part of human clinical trials^29^(NCT03300050).

This strategy of chimeric immunization has been extended to SARS-CoV-2 by Baric and co-workers^30^. In this work, mice were immunized with different chimeric sarbecovirus S proteins. The best-performing chimeras and immunization strategies were found to elicit antibodies with superior breadth of protection when compared to standard immunization regimens when assessed for protection against related sarbecoviruses in both neutralization assays and challenge studies.

Based on these successes, we aim to develop a systematic approach for designing chimeric coronavirus antigens. To accomplish this, we leverage the recently-developed AlphaFold structure-prediction method and assess its use in stability prediction for chimeric protein design. AlphaFold2 is a powerful, neural-network-based protein structure prediction algorithm that substantially increased prediction accuracy over prior methods^31^ and has been rapidly adopted for different protein design applications^32-34^, also spawning a set of related deep-learning approaches for protein structure prediction^35,36^. AlphaFold 3^37^ of course offers additional algorithmic changes, but version 2 was used for this work in part due to the modeling throughput required. This approach complements prior methods for stability prediction, both those that utilize global structure and those that focus on point mutations^38-43^. One advantage of AlphaFold over prior methods is that it can in a single step produce predictions of both structure and global stability, particularly for large domain swaps.

We report both a pipeline and results for chimeric antigen design, where chimeras are formed by replacing the S1 domain of the SARS-CoV-2 S protein with the S1 domain from another coronavirus spike protein, leaving the S2 domain conserved across all chimeras (Figure 1). This approach complements computational designs focused on yielding stable S2-only immunogens^44^. We compare results against molecular dynamics simulation and experimental expression and stability testing and present a set of top-ranked antigens that will be used for immunological testing.

**Figure 1.**
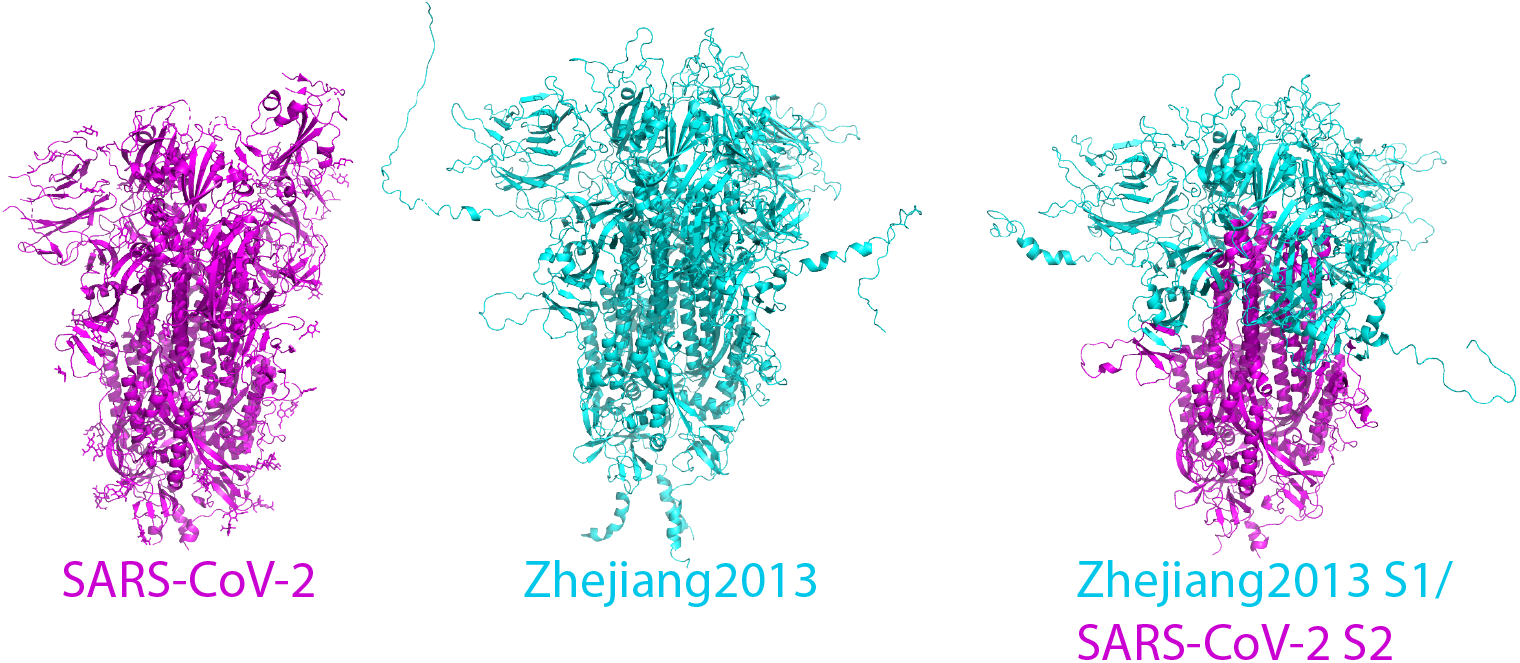
Design of chimeric spike proteins. The process of chimeric S1/S2 spike protein design is illustrated for a sample betacoronavirus sequence. The sequence of SARS-CoV-2 (structure rendered in magenta) is spliced with that of Bat Hp-betacoronavirus/Zhejiang2013 (predicted structure rendered in cyan) at the predicted S1/S2 junction. The predicted structure of the chimera is rendered in a mix of cyan and magenta representing the parental sequences for each domain. pLDDT (predicted local distance difference test) scores from AlphaFold for both the parental domains and chimeric sequences are calculated, and these are averaged per residue to yield the relative stability score.

## Results

To facilitate prediction of stable chimeras and evaluation of the results, we generated a curated set of 115 coronavirus spike protein sequences (Table S1). These sequences are evolutionarily diverse and display particular variability in the receptor-binding and N-terminal domains, as demonstrated in an analysis of sequence entropy rendered on the SARS-CoV-2 S protein structure (Figure 2). Domain boundaries were identified via sequence alignment, and chimeras were generated computationally by splicing the S1 domain from each of the 114 other sequences onto the S2 domain of SARS-CoV-2 (Figure 1). For comparison, a similar operation was performed to splice RBDs from each of the 114 other chimeras onto SARS-CoV-2; those results are presented in the Supplement (Figs. S1-S2, Table S3). Structures for each of these chimeras were then predicted using AlphaFold2, and the corresponding local confidence scores (pLDDT) recorded for use in stability predictions.

**Figure 2.**
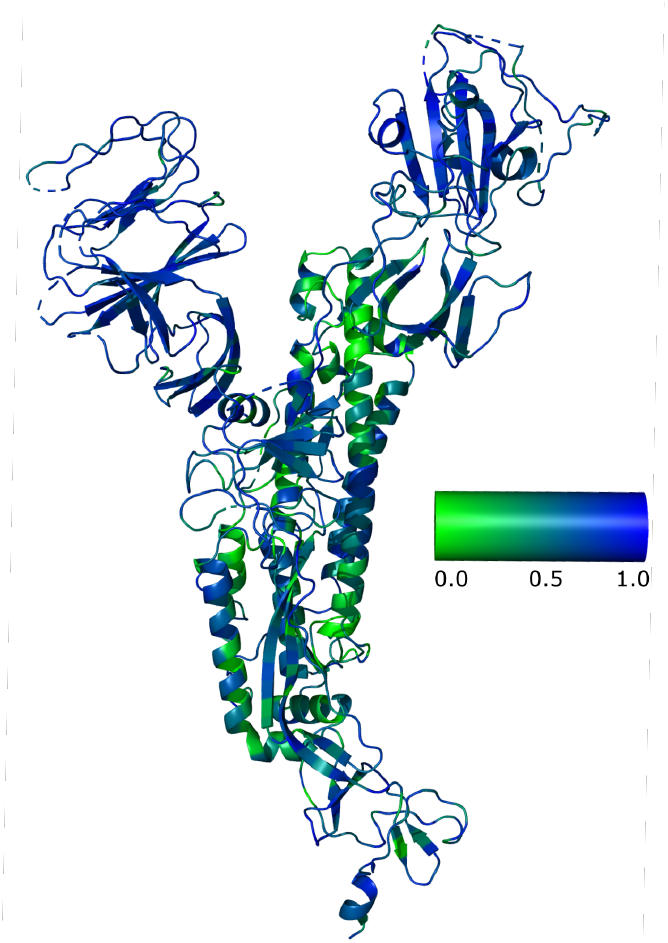
Relative sequence entropy among analyzed coronavirus spike proteins. Sequence entropy was calculated for a multiple sequence alignment of all coronavirus sequences used and rendered on the 6VSB structure, where green represents lowest sequence entropy and blue represents highest. As expected, the receptor-binding domain and N-terminal domain have the greatest sequence entropy, and loops tend to have higher entropy than adjacent structural elements.

Because each of the 114 spike protein sequences other than SARS-CoV-2 also corresponds to a native “wild-type” virus, it must represent a stable folded protein. We therefore evaluated the predicted gain or loss of stability for each of the 114 spliced S1 domains in its chimera with SARS-CoV-2 versus in its full-length native sequence. The analogous comparison was also performed for the region of the chimera derived from SARS-CoV-2 (the S2 domain). Stability scores were thus defined for each chimeric sequence *i* generated by splicing coronavirus *i* and SARS-CoV-2 as:

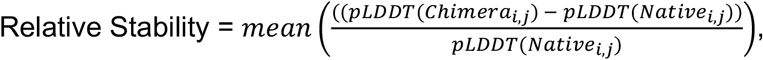

where pLDDT are predicted local distance difference test^45^ scores emitted by AlphaFold, Native_*i,j*_ was defined as the residue corresponding to *j* in either SARS-CoV-2 or coronavirus *i*, depending on which native protein the residue derived from.

Relative stabilities were calculated according to the above relationship and plotted versus sequence similarity in Figure 3. As expected, coronavirus S1 sequences highly similar to SARS-CoV-2 tended to yield stable chimeras, while sequences less similar to SARS-CoV-2 tended to yield less-stable chimeras. For comparison, the HexaPro stabilized SARS-CoV-2 spike construct^46^ was also modeled and yielded a relative stability of -8.0%. This makes sense because the protease-resistance mutations are not reflected in AlphaFold scoring, so the predicted stability increase mostly reflects stabilization of flexible regions. The Spearman rank correlation between sequence similarity and relative stability was calculated at 0.51 for the entire group of chimeras. The chimeras of greatest potential utility, however, were those that *violated* this sequence-stability correlation: where a sequence highly divergent from SARS-CoV-2 yielded a chimera with minimal decrease in stability. If correct, these would represent stable proteins that differ antigenically from SARS-CoV-2 in the exchanged region. For comparison, we also computed FoldX^38^ stability scores for all chimeras; these are plotted in Fig. S3 and show a 0.89 Spearman rank correlation with pLDDT and a 0.78 Spearman rank correlation with relative stability. Overall, the predicted ranking of chimeras is similar, but there are some substantial outliers.

**Figure 3.**
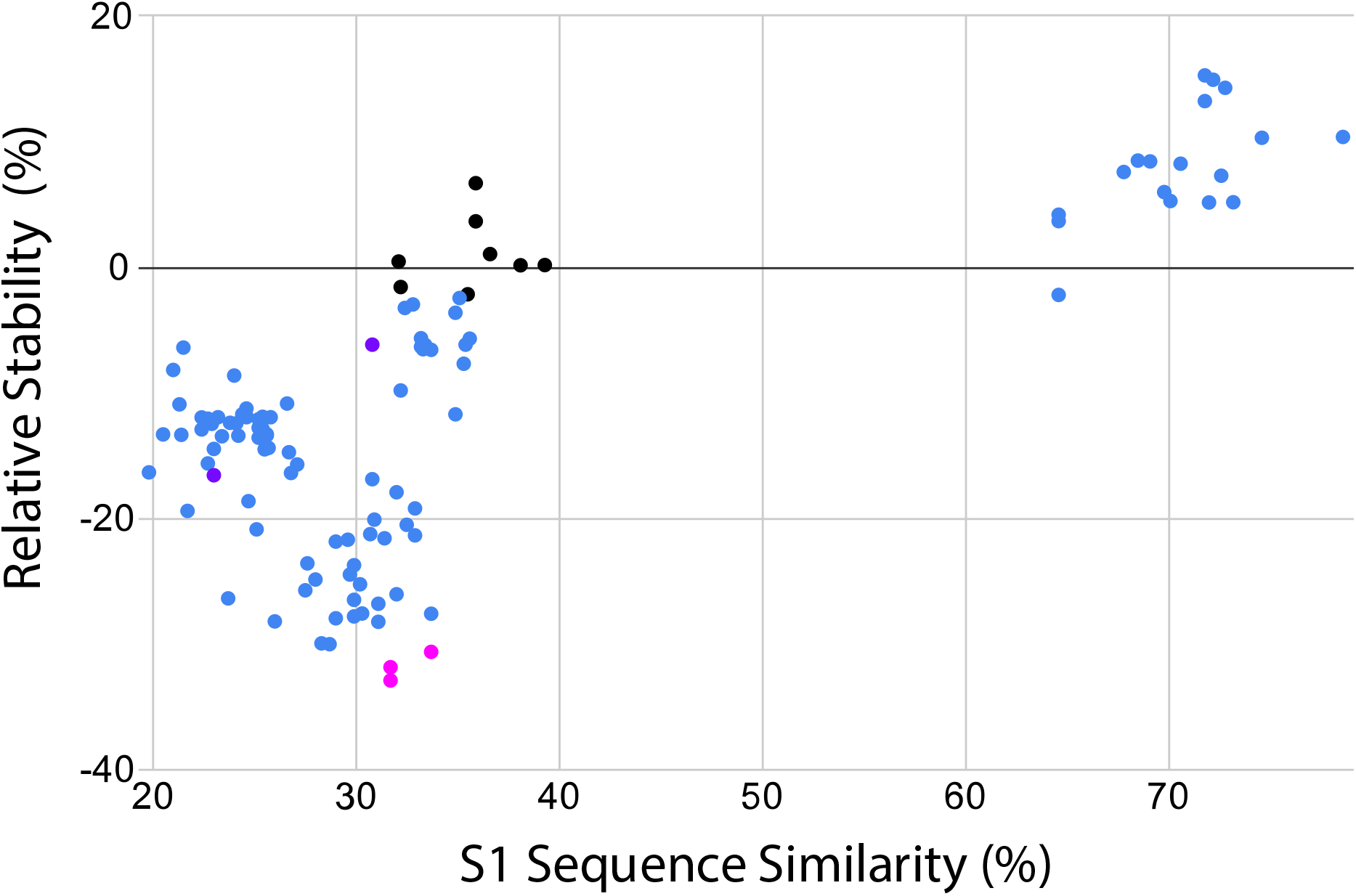
Predicted stability of coronavirus spike chimeras. The relative stability score for each S1/S2 chimera is plotted against the sequence similarity between the parental S1 sequence and SARS-CoV-2. The similarity between the two domain sequences was determined using EMBOSS^47^. Sequences fall into two broad groups: one with high similarity and a gain in relative stability, and one with low similarity and a loss in relative stability. The outliers that are low similarity but relatively high relative stability are of particular interest for immunogen design. Predicted high-stability chimeras selected for simulation are plotted in black, and predicted low relative-stability chimeras used as controls are plotted in magenta. Additional low overall-stability controls are plotted in purple. Data are tabulated in Table S4.

The sequences in our analysis showed a bimodal similarity to SARS-CoV-2, with most sequences ≤40% similarity or ≥65% similarity in the S1 domain. Because our goal was to identify antigenically different chimeras, we focused on the low-similarity group and selected the eight chimeras in this group with the highest relative stability for further analysis. Each of these chimeras was simulated using molecular dynamics for 100 ns as a computational proxy for physical stability. Structures were evaluated for unfolding as well as structural fluctuations, and the results are plotted in Figures 4-6. Five low-similarity, low-stability sequences were also simulated as negative controls (three based on low relative stability and two on low mean pLDDT).

**Figure 4.**
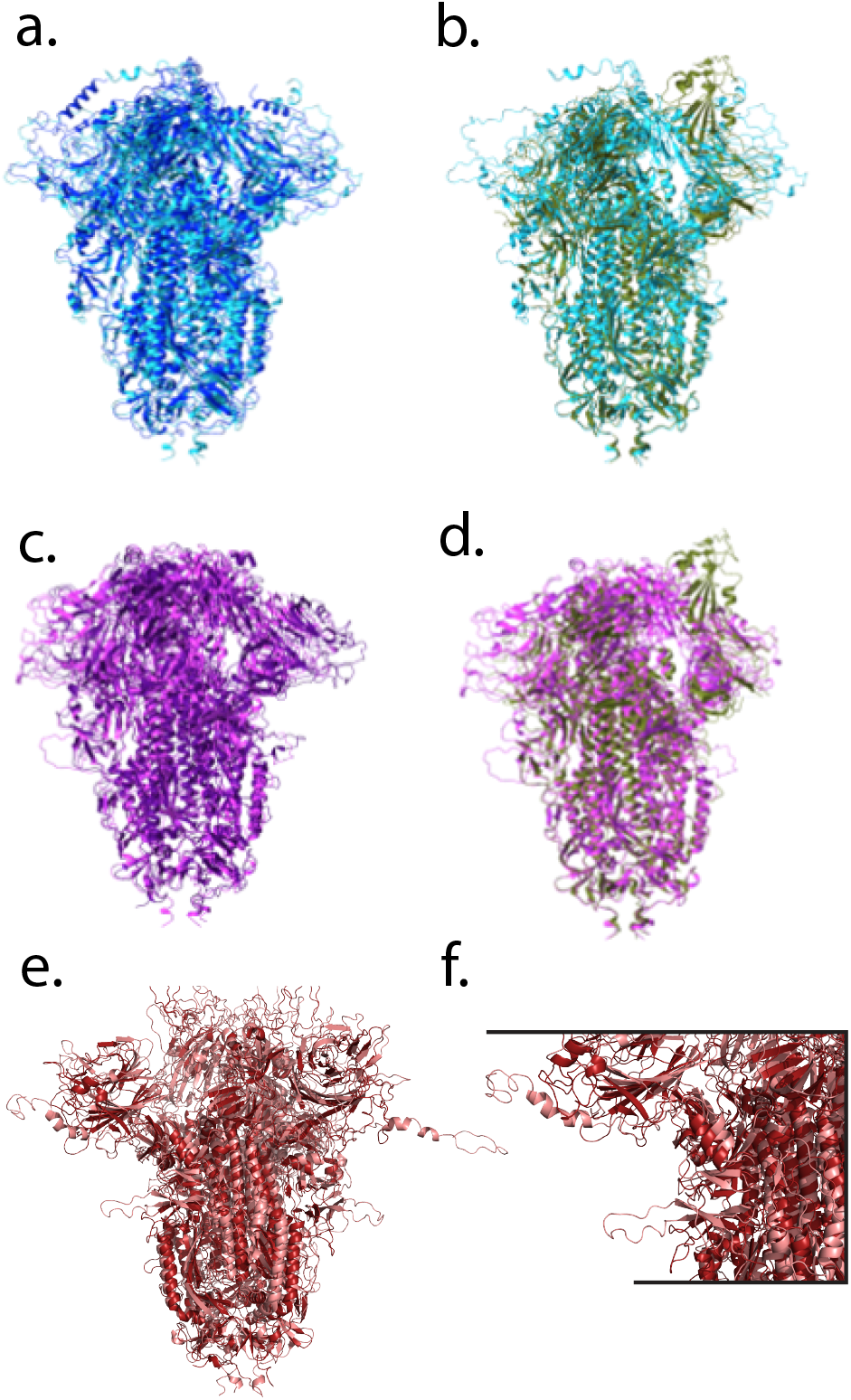
Predicted and simulated structures of top-scoring chimeras. Structures of two of the top-scoring chimeras are rendered as follows. Chimeras from Rousettus bat coronavirus GCCDC1 (panels (a) and (b)) and Hedgehog coronavirus 1 (panels (c) and (d)) are shown before and after 100 ns of molecular dynamics simulation (left column), with the before and after structures also superimposed on renderings of the SARS-CoV-2 6VSB PDB structure (panels b and d, dark green). The chimera from Bat Hp-betacoronavirus Zhejiang2013 is rendered in panel (e), with a zoomed rendering in panel (f) showing the n-terminal and furin-cleavage loops that were initially modeled as extended becoming more compact over the simulation. For each chimera, the structure before simulation is rendered in a lighter shade and after simulation is rendered in a darker shade. The 6VSB SARS-CoV-2 structure has one receptor-binding domain in the “up” conformation, whereas both chimeras rendered are modeled as fully down and remain so throughout the simulation.

All predicted-stable chimeric proteins simulated showed minor structural fluctuations in the first 5 ns of simulation and then remained stable for the remainder of the simulation (Fig. 5). Structural changes were limited; some loops and termini that were initially modeled as disordered became more compact and acquired secondary structure (Fig. 4f). These changes for Zhejiang2013 caused the root-mean-squared deviation (RMSD) to be slightly higher than that of the lowest-RMSD control chimera, HKU15). Much of the conformational fluctuation observed in the simulations was concentrated near the furin cleavage site and similar loops; as well as the N-terminus and C-terminus (Figure 6). In part because the S1/S2 loop was disordered in the original full-length SARS-CoV-2 structures and mutated away in others ^6,48^, it is unsurprising that AlphaFold modeled this loop in a physically less plausible state. In chimeras expressed as stable immunogens, we would expect to follow a strategy similar to that pursued successfully with SARS-CoV-2 ^46^ where either the furin site was mutated or the S1/S2 loop truncated altogether. Furthermore, the S2 domains for the predicted-stable chimeras remained highly stable throughout the simulations (Fig. 6), as would be hoped for constructs designed to help focus an immune response against conserved S2 epitopes. Predicted-unstable chimeras showed much more structural fluctuation. This was due to a combination of two factors: regions that were intrinsically less stable (correct AlphaFold predictions; Fig. S4) and regions that became much more compact over the simulation (potentially incorrect AlphaFold predictions). This suggests that the scoring method used may have some false-negative results – chimeras where the predicted stability is much less than the simulated stability – but few or no false-positive results – predicted-stable chimeras were indeed highly stable on simulation.

**Figure 5.**
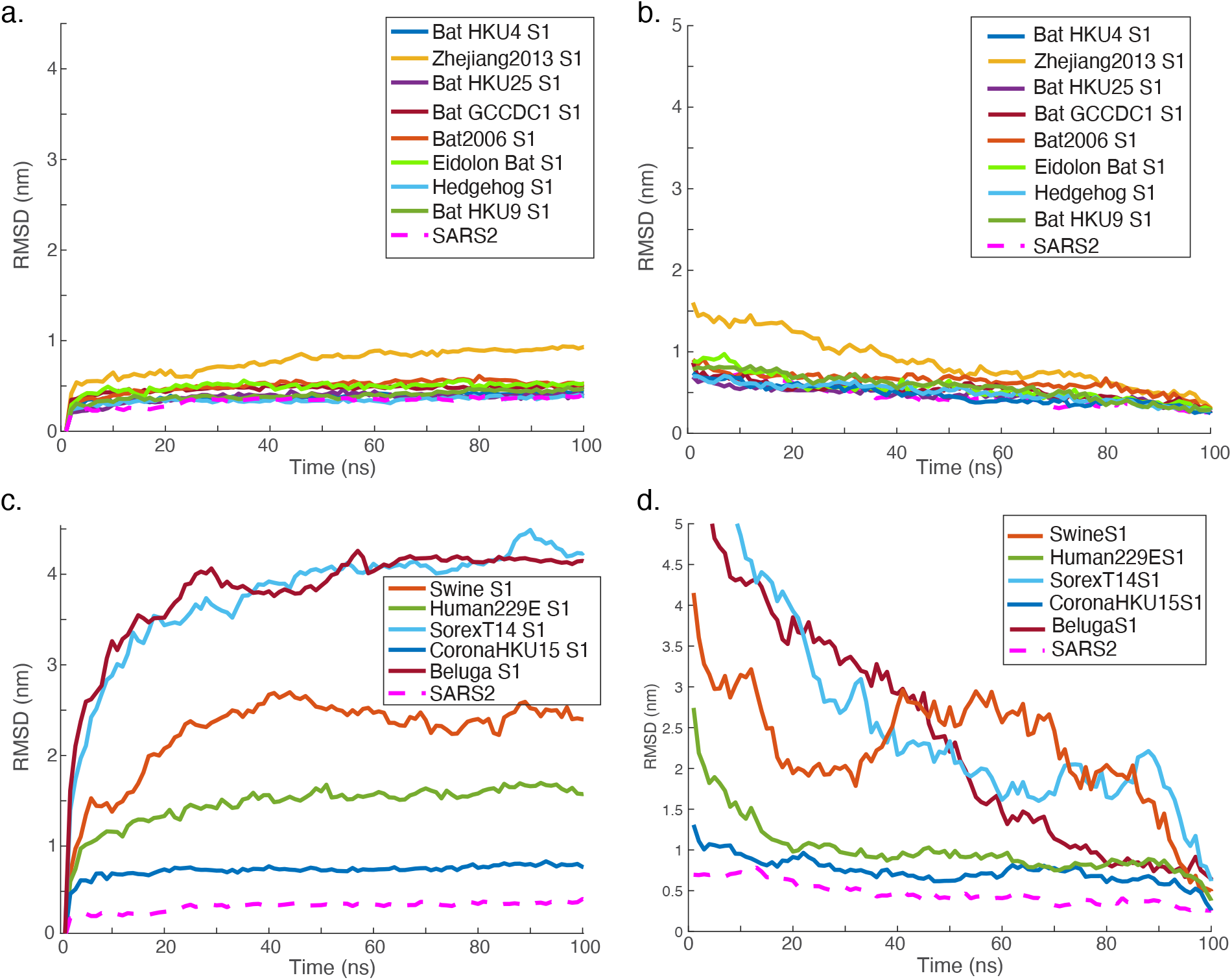
Structural stability of simulated chimeras. Molecular dynamics simulations were computed for predicted structures and root-mean-squared deviation (RMSD) from a reference structure is plotted versus time. Plots are given for (a-b) the eight chimeras in the low-sequence-similarity cluster with the highest predicted relative stability, (c-d) five low-stability controls. SARS-Cov-2 is plotted in both panels as an additional control. Panels (a,c) plot RMSD from the starting structure, whereas panels (b,d) plot RMSD from the ending structure. As expected, the low-stability controls showed the greatest RMSD, with one showing gross structural changes. The eight predicted-stable chimeras and SARS-CoV-2 all showed an initial increase in RMSD followed by a stabilization over the course of the 100-ns simulation. This can also be seen in RMSD plots relative to the end of the simulation (Fig. S5).

**Figure 6.**
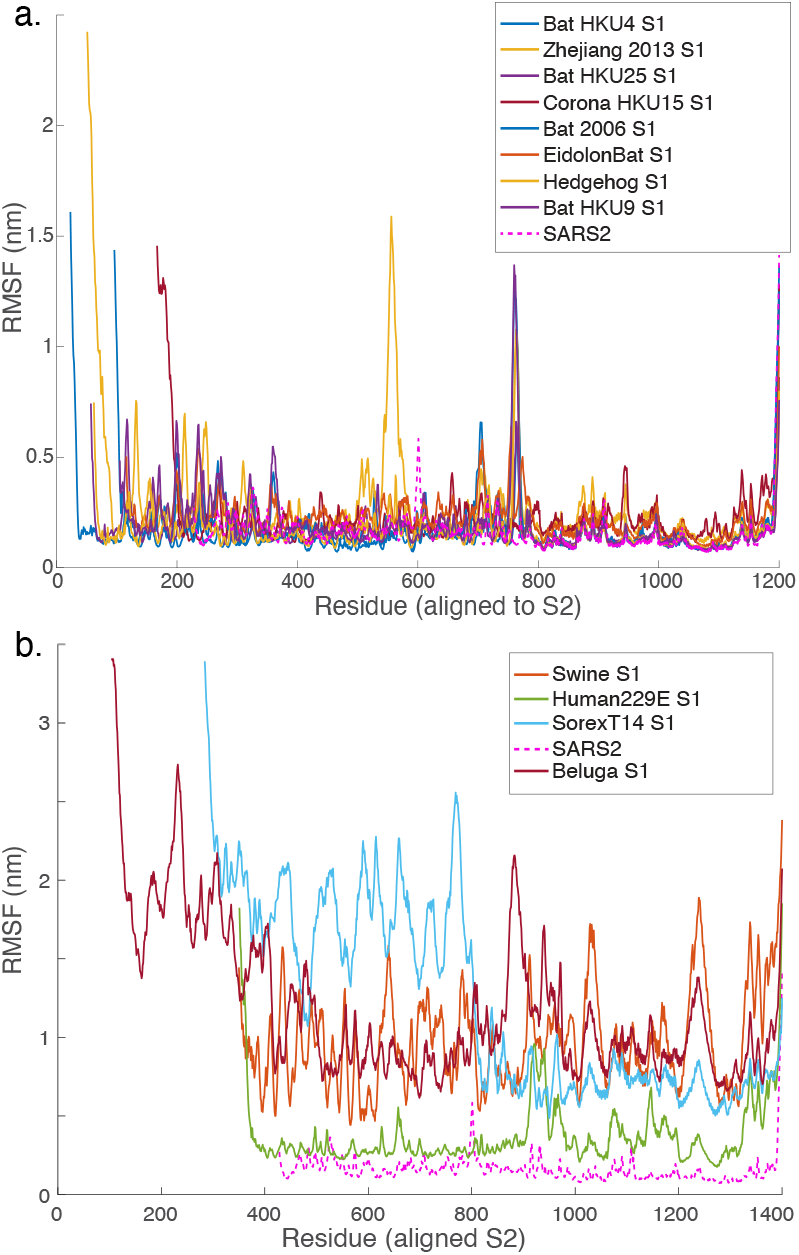
Per-residue structural stability of simulated chimeras. Root-mean-squared fluctuation (RMSF) values are plotted per residue for each of the simulated chimeras. Values were calculated on nanosecond intervals throughout the simulation trajectory. Panel (a) shows the predicted high-stability chimeras, and panel (b) shows the predicted low-stability chimeras. SARS-CoV-2 is included as a comparator. As expected, major loops as well as the C- and N-termini show the greatest fluctuation in the high-stability chimeras, and the low-stability chimeras have globally greater fluctuations.

Experimental testing was performed by expressing chimeric full-length spike proteins on pseudoviral particles in mammalian cell culture, purifying the particles, and assaying stability. Five predicted-stable chimeras were tested, along with two predicted-unstable chimeras and native SARS-CoV-2 as a comparator. Similar to previous work^46^, expression of spike protein was assayed as a first indicator of stability. In this case, expression was measured via spike anti-S2 immunofluorescence on individual pseudoviral particles. The results (Fig. 7) show substantially stronger expression for Eidolon Bat S1 and Bat GCCDC1 S1 chimeras than for native SARS-CoV-2, while Bat2006 S1 yielded similar expression on purified pseudoviral particles, and Zhejiang2013 S1 yielded somewhat lower expression. The two predicted-unstable chimeras, Swine S1 and Sorex T14 S1, yielded 2.5-fold to 12-fold lower, along with one predicted-stable spike, Bat HKU25 S1, at 4-fold lower expression, which we regard as a negative result. Of note, measuring expression via immunofluorescence also demonstrates that major S2 epitopes are stably displayed on the surface of pseudoviral particles expressing the chimeras and capable of binding antibody.

**Figure 7.**
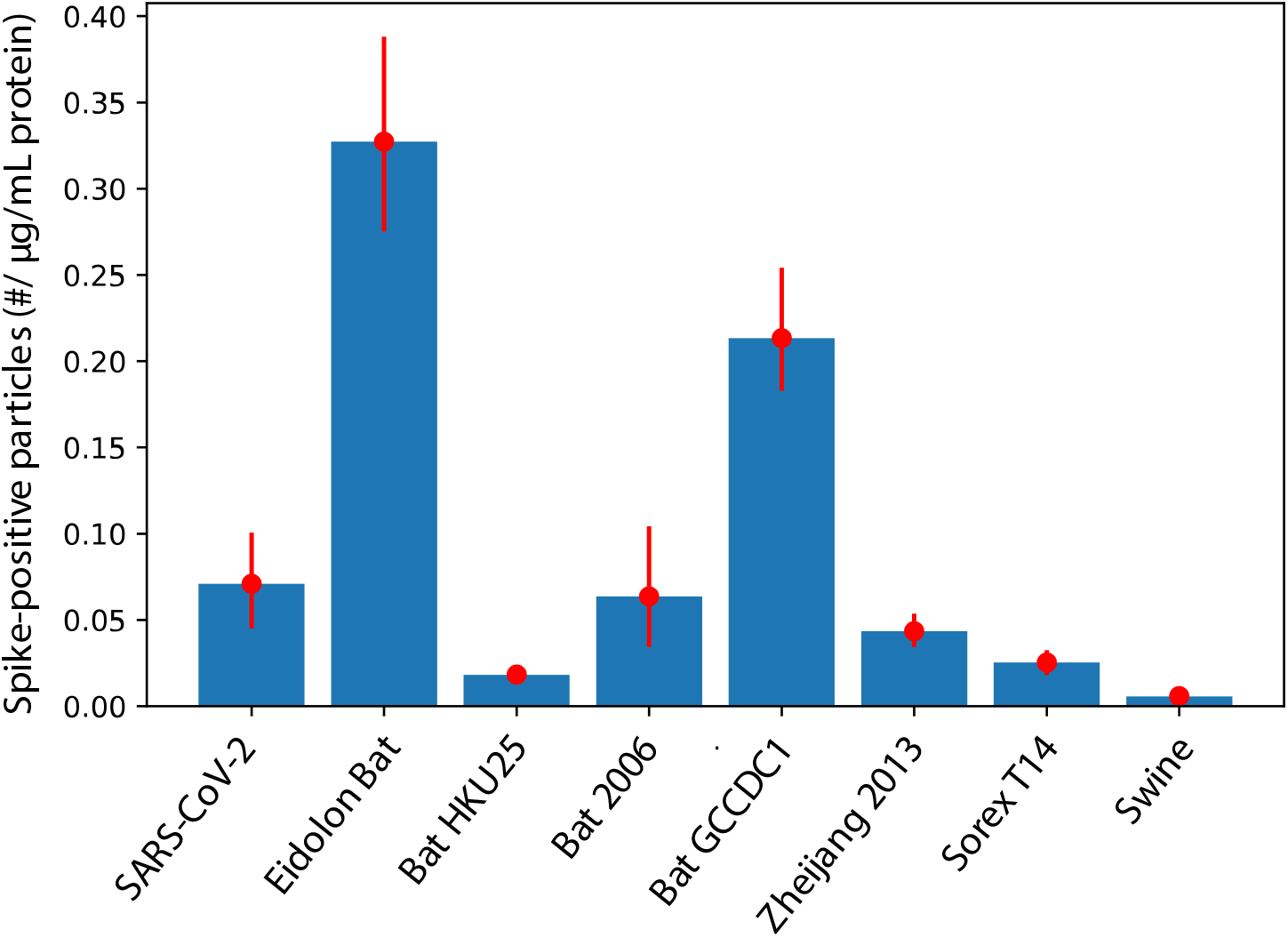
Expression of chimeric spike proteins on purified pseudoviral particles as assessed by single-particle immunostaining. Pseudoviral particles were purified, biotinylated, immobilized in a microfluidic flow cell via a biotin-neutravidin-biotin linkage, and then immunoassayed using an anti-S2 antibody. The number of spike-positive particles was then scaled by the total protein in the pseudoviral preparation to give a relative measure of spike expression per pseudovirus, which is plotted here. Eidolon Bat and Bat GCCDC1 showed substantially better expression than native SARS-CoV-2, Bat2006 was comparable to native SARS-CoV-2, while Zhejiang2013 was somewhat lower. BatHKU25 and the two predicted-unstable chimeras, Swine and Sorex T14 showed substantially worse expression, likely indicating spike instability at the stage of expression and pseudoviral budding. Error bars indicate 90% confidence intervals from bootstrap resampling across fields of view.

Further testing of thermostability was performed using differential scanning fluorimetry of pseudoviral particles using particles lacking any spike construct as a negative control. All thermostability assays of course rely upon sufficient levels of expressed spike protein, so low-expressors are expected to behave more like “bald” pseudoviral particles in these assays.

Thermal profiles are shown in Fig. 8. SARS-CoV-2, Eidolon Bat S1, and Bat2006 S1 all show a bimodal unfolding profile with inflection points in the 50-60 ºC range. Surprisingly for the degree of protein expression observed, Bat GCCDC1 S1 did not show a high-magnitude unfolding peak over that temperature range. However, it did display substantial shoulders on the underlying pseudovirus particle unfolding peak that register as deviations from bald pseudovirus (Figs. 8b-c) and may represent higher-temperature unfolding than native SARS-CoV-2, in the 70-80 ºC range. Definitive assignment would require further confirmation, but the overall profile is consistent with a stable chimeric spike, in accordance with prediction. Overall, these sets of experiments suggest that 3/5 predicted-stable chimeras were indeed equivalently stable to native SARS-CoV-2, one was intermediate, and one was relatively unstable. 2/2 predicted-unstable chimeras were indeed unstable upon expression at 37 ºC in mammalian cell culture.

**Figure 8.**
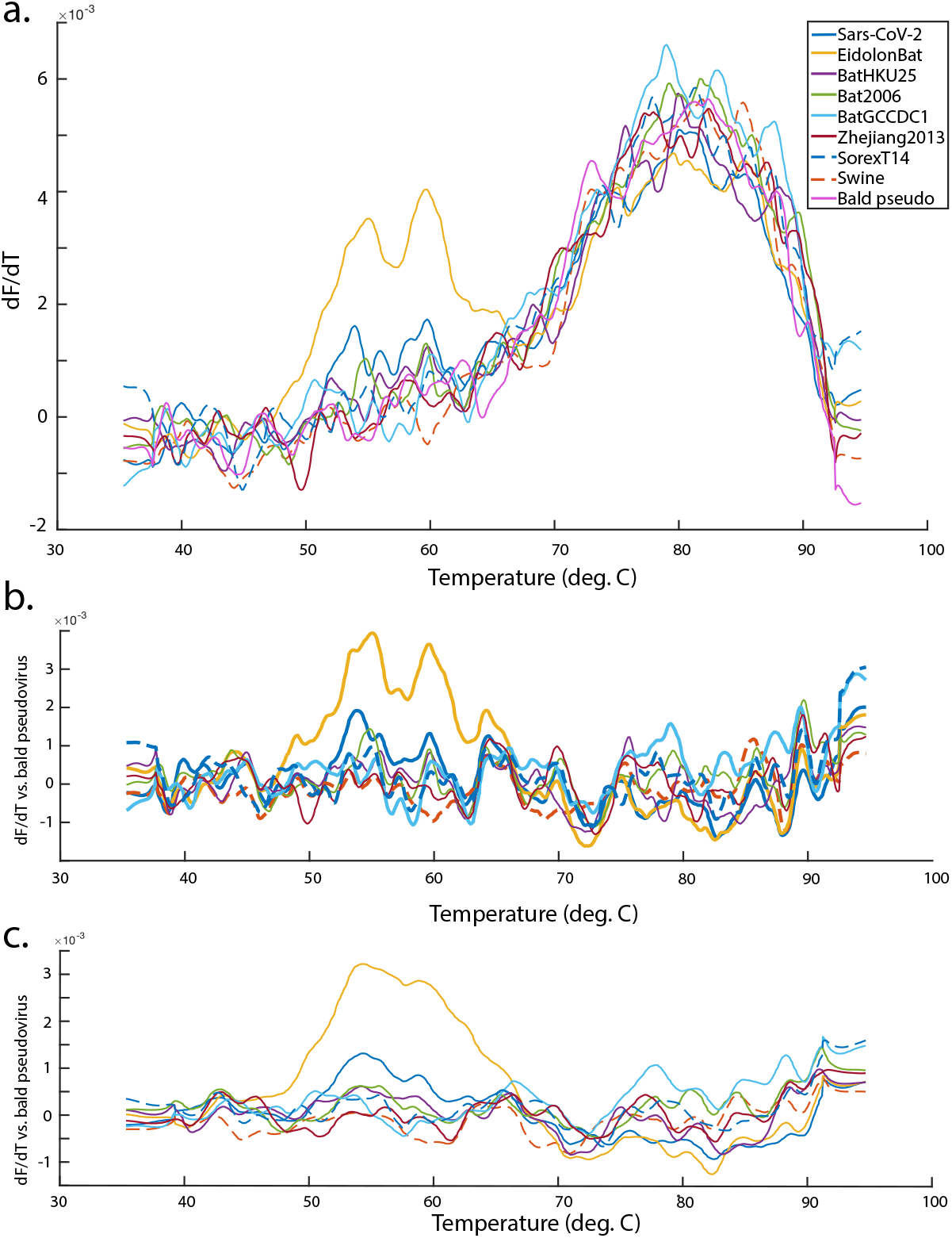
Thermal stability of pseudoviruses expressing chimeric spike proteins. Plotted are dF/dT values for pseudoviruses expressing SARS-CoV-2, 5 predicted-stable chimeric spikes, 2 predicted-unstable chimeric spikes, and bald pseudovirus produced with no spike. The dF/dT values (first derivative of the 350/330 nm fluorescence ratio with respect to temperature) are plotted in panel a, and the background-subtracted dF/dT are plotted in panels b-c, using bald pseudovirus as an estimator of non-spike background. Fluorescence ratios were smoothed using locally weighted scatterplot smoothing and showed peaks that were robust to choice of smoothing parameter over the range 40-80, with smoothing parameter of 50 shown in a-b and 80 shown in c. Eidolon bat chimeras showed stability comparable to native SARS-CoV-2, while Bat GCCDC1 chimeras were less well determined but consistent with thermal stabilization relative to SARS-CoV-2. Bat 2006 chimeras showed lower signal but were likely similar to SARS-CoV-2. Predicted-unstable chimeras had poor expression on pseudovirus produced in cell culture at 37 C (Fig. 7) and were thus likely unstable at the point of expression and pseudoviral budding.

## Discussion

Here, we introduce a measure of relative stability to assess chimeric sequences: the predicted stability relative to the parent sequences for each portion of the chimera. As assessed by molecular dynamics simulations of the AlphaFold models, relative stability is a fairly effective metric for selecting stable chimeras. An alternate metric, however, would be average pLDDT, or absolute predicted stability. These two metrics rank the chimeras examined here very similarly, with a Spearman rank correlation of 0.86. However, the ranking within the top 8 chimeras selected for simulation is essentially uncorrelated (Spearman rho 0.095). Relative stability is a much stronger prediction of stability in molecular dynamics simulation (Spearman rho 0.38 vs. -0.048 with average RMSD over each 100 ns simulation), although both of these are imprecise predictors of the molecular dynamics simulations. Our results thus suggest that relative stability is a better scoring metric for chimeric proteins but that a secondary screen with molecular dynamics simulation adds substantial further information. Experimental testing is largely concordant with the computational predictions, with the stable chimeric spikes showing long-term stability in molecular dynamics simulations, the predicted-unstable spikes being unstable throughout, and one of the two experimentally least-stable among the original positive predictions showing intermediate stability in molecular dynamics simulations. The only true outlier was Bat HKU25 S1, which showed good stability in molecular dynamics simulations yet expressed poorly.

It is interesting to speculate as to why absolute stability may perform less well than relative stability. Relative stability scores take the native viral spike proteins as a baseline and measure predicted stability compared to those. They thus will correct for any AlphaFold scoring errors on the native sequence, but conversely they will fail to capture variation in stability among native viral spike proteins. Thus, on the dataset examined here, AlphaFold prediction errors appear to be somewhat greater than variation in native viral spike stability, at least as scored by molecular dynamics simulation for the most stable predicted sequences.

AlphaFold may have some generality as a predictor of protein stability for protein-design applications; here we use it specifically to predict chimeras of otherwise well-structured proteins that have a fair degree of overall homology. We use AlphaFold 2 because of its source-code availability (and it was the current version at time of project initiation), but it will be informative to systematically compare results with AlphaFold 3^49^. Within this more limited domain, the relative stability metric shows good success in guiding selection of chimeras that are stable as assessed by molecular dynamics simulation. Ultimately, experimental expression and testing of chimeric proteins will be required to test the best candidates. It is hoped that this approach of AlphaFold prediction, molecular dynamics simulation, followed by experimental testing and optimization as needed will assist in the design of chimeric immunogens against betacoronaviruses as well as other viral pathogens.

## Methods

### Sequence curation

Sequences were downloaded from the NCBI Virus portal on 10/1/2021 using the following search criteria: Viruses: Alphacoronavirus, Betacoronavirus, Gammacoronavirus, Deltacoronavirus; Proteins: spike, spike protein, spike glycoprotein, surface glycoprotein, spike surface glycoprotein, membrane glycoprotein; Sequence length greater than 700 and less than 2000. All duplicate sequences and SARS-CoV-2 sequences were removed. The resulting sequences are given in Table S1. The reference sequence for SARS-CoV-2 was defined as the residues that were structurally resolved in chain B of the 6VSB PDB structure^50^.

### Sequence entropy calculation

All analyzed coronavirus spike sequences were aligned to the SARS-CoV-2 reference sequence, and substitution scores were calculated for each position using the BLOSUM62 matrix. Sequence entropy was then calculated according to the equation

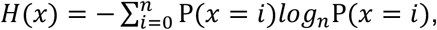

where P*(x=i)* is the probability of BLOSUM value *i* at position *x*, and *n* is the number of possible BLOSUM values.

### Chimera Sequence Generation

The S1 domain boundary was taken as previously defined^51^ and applied to the SARS-CoV-2 reference sequence as defined above. Pairwise sequence alignments were created between SARS-CoV-2 and each other spike protein sequence retrieved above using Muscle^52^ (v.3.8.31), and this alignment was used to define S1 domain boundaries across the spike proteins examined. Sequence similarity between S1 domains was scored using EMBOSS^47^ (v6.6.0, alignment using the Needleman-Wunsch algorithm). Custom Python code (available from https://github.com/kassonlab/coronavirus-chimera-prediction) was used to generate 114 chimeras swapping the S1 regions from each of the other coronaviruses with the S1 of the SARS-CoV-2 reference sequence.

### AlphaFold Modeling

The multimer prediction algorithm in AlphaFold^31,53^ (v.2.2.2) was used to analyze the 115 wild-type spike proteins (SARS + 114 others) and 114 chimeric spike proteins. An additional 10 sequences did not yield AlphaFold predictions and are listed in Table S2; these were excluded from subsequent analysis. Databases used were reduced BFD for HHblits, PDB seqres for hmmsearch, and UniProt for JackHMMer. Max template date was set to 09-12-2022, and 5 models were created per sequence with the top-scoring model selected after relaxation. Calculations were performed on NVIDIA A100 GPUs, enabling unified memory in Tensorflow to facilitate modeling of long trimeric sequences.

### Relative Stability Scores

Relative stability scores were calculated as follows for each residue *i*:

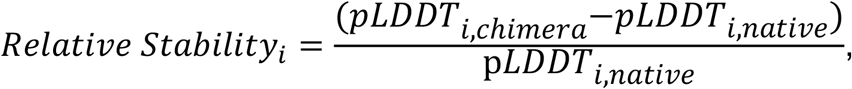

where *pLDDT*_, chimera_ is the pLDDT calculated for residue *i* in the chimeric sequence and *pLDDT*_*i, native*_ is defined as the pLDDT calculated for residue *i* in the context of the full sequence from which that amino acid derived (in the case of S1/S2 chimeras, SARS-CoV-2 for S2 and the variable coronavirus sequence for S1). Per-residue relative stability scores thus calculated were averaged across the full trimeric protein to yield a single score for each chimera.

### Molecular Dynamics Simulations

Molecular dynamics simulations were run using GROMACS^54^ (v2021.2) for each of the 8 chimeras with the greatest relative stability and sequence similarity <40%. Simulations were performed using the CHARMM36 force field^55^. Each protein was placed in an octahedral periodic box with a minimum of 1 nm distance to the box boundary and solvated with TIP3P water and 150 mM NaCl. Simulations were run using hydrogen bonds constrained with LINCS and long-range electrostatics treated with Particle Mesh Ewald, temperature set to 310K with the velocity-rescaling thermostat^56^, and pressure maintained at 1 bar with the Parrinello-Rahman barostat. Simulations were run for 100 ns using 2-fs timesteps.

### Creation of chimeric protein expression vectors

S1 domains corresponding to five predicted-stable chimeras and two predicted-unstable chimeras were synthesized from sequence by Twist Biosciences. Full-length SARS-CoV-2 Wuhan spike with a C-terminal ALAYT mutation was a kind gift of Jesse Bloom and used in his and our previous work^57,58^. Each S1 was amplified via PCR and Dpn1 digested, as was the native SARS-CoV-2 expression plasmid minus SARS-CoV-2 S1. The digested S1 product (range 50-150 ng) was mixed with S2-vector product in a 1:2 ratio, added to 10 µl of 2x Gibson Assembly master mix (New England Biolabs), volume-adjusted to 20 µL with water, and incubated at 50 ºC for 45 minutes. Transformation was performed as follows: 40 µl of XL10 Gold high efficiency *E. coli* (Agilent) was thawed for 3 minutes and mixed with 2 µl of Gibson Assembly product and incubated for 30 minutes on ice. Each tube was heated to 42 ºC for 30 seconds and placed on ice for 2 minutes after adding 350 µl of SOC media. Tubes were shaken at 190 rpm and 37°C for 1 hr, then plated on a 1 mg/ml Amp agar plate to incubate overnight at 37 ºC. A single colony was selected for an overnight 1 mg/ml LB broth culture. The plasmids were extracted, verified via long read sequencing, and stored for pseudovirus expression.

### Pseudovirus expression and purification

Chimeric spikes were purified and expressed in a variant of a previously published protocol ^58^. The following plasmids were mixed in a tube: 0.85 µg chimeric spike plasmid, 2.5 µg luciferase-IRES-ZsGreen (BEI NR-52516), 0.55 µg HDM-Hgpm2 (BEI NR-52517), 1.62 µg pRC-CMV-Rev1b (BEI NR-52519), and 0.55 µg HDM-tat1b (BEI NR-52518. Opti-MEM media was added for a total volume of 236 µL, and 14 µl of lipofectamine was added to 236 µl of OMEM and incubated for 5 minutes. This mixture was used to transfect 293T cells in a 75-cm^2^ flask; media was replaced after 6 hours, and supernatant harvested after an additional 48 hours. Supernatant was centrifuged at 800 rcf, 4 ºC for 8 min and passed through a 0.45 µm filter to remove cell debris. Pseudoviral particles were crudely purified by ultracentrifugation at 140,000 rcf, 4 ºC for 2 hours on a 2.5 ml 25% sucrose/HEPES cushion. Pellets were resuspended in HEPES-MES buffer and rocked gently overnight at 4 ºC.

### Immunofluorescence

Pseudoviral particles were fluorescently labeled as previously described^57^ by incubation with 1% Texas Red-DHPE in HEPES-MES 2 hours at room temperature with gentle rocking, followed by overnight incubation with biotin-PE (83 nM final concentration). A 2-µL aliquot was diluted 1:3 in HEPES-MES and added to a microfluidic flow cell functionalized with PLL-PEG-Biotin-neutravidin as previously described^57^. The mixture was incubated for one hour at room temperature, washed twice with 500 µL of HEPES-MES, and blocked with 10 µL of 2% normal goat serum, and washed once more. Primary immunolabeling was performed with 10 µl of anti-S2 monoclonal antibody (Invitrogen MA5-35946) at 1 µg/ml, incubated overnight at 4°C, then washed twice. Secondary immunolabeling was performed using an Alexa 488-labeled goat anti-mouse antibody (Invitrogen) at 1 µg/ml with a 45-minute incubation followed by a 45-minute incubation and two washes. Imaging was performed on a Zeiss AxioObserver A2 microscope with a Lumencor Spectra-X light engine and an Andor Zyla 4.2plus sCMOS camera, with measurements recorded on five fields of view per condition. Background-subtracted fluorescence intensity was used as a measure of relative spike concentration, controlled against total pseudoviral protein from a microBCA assay.

### Nanoscale differential scanning fluorimetry

Pseudoviral particles were diluted to 30 µg/ml of total protein as per microBCA performed above. Fluorescence was measured in a Nanotemper Tycho NT.6, using a scanning rate of 30 °C/minute from 35 °C to 95 °C. The ratio of fluorescence between 350 and 330 nm was measured over this temperature range and smoothed using locally weighted scatterplot smoothing. To measure robustness to smoothing, the smoothing parameter was varied between 40 and 80, and the conclusions drawn remained unchanged over this range. The derivative of fluorescence ratio with respect to temperature (dF/dT) was measured and used to assess thermal unfolding. Particle size distributions were additionally characterized by nanoparticle tracking analysis at both room temperature and after heating to 55 ºC. The primary populations were all consistent with monomeric pseudovirus, and there was not a major difference in aggregate formation between SARS-CoV-2 expressing pseudoviruses and bald pseudoviruses after heating, suggesting that, in these expression regimes, spike unfolding is not a major driver of pseudovirus aggregation.

## Code and data availability

Code and scripts used to generate the chimeras, run AlphaFold, and analyze resulting scores are available from https://github.com/kassonlab/coronavirus-chimera-prediction. Multiple sequence alignments used are also in the above GitHub repository. Predicted structures simulated and analyzed as well as primary experimental data are available from the Zenodo repository doi: 10.5281/zenodo.13830740.

## Acknowledgements

This work was supported by NIH R01 GM138444, a Coulter Translational Research Award, and a Wallenberg Academy Fellowship to P.M.K. J.S. was also supported by NSF 1810762. Computational resources were provided by Research Computing at the University of Virginia and the PDC Center for High-Performance Computing supported by the Swedish National Infrastructure for Computing.

## References

1. Organization, W.H. WHO Coronavirus dashboard. Vol. 2024 WHO Coronavirus (COVID-19) Dashboard (2024).

2. Jackson, C.B., Farzan, M., Chen, B. & Choe, H. Mechanisms of SARS-CoV-2 entry into cells. Nat Rev Mol Cell Biol 23, 3–20 (2022).

3. Hoffmann, M. et al. SARS-CoV-2 Cell Entry Depends on ACE2 and TMPRSS2 and Is Blocked by a Clinically Proven Protease Inhibitor. Cell 181, 271–280 e8 (2020).

4. Letko, M., Marzi, A. & Munster, V. Functional assessment of cell entry and receptor usage for SARS-CoV-2 and other lineage B betacoronaviruses. Nat Microbiol 5, 562–569 (2020).

5. Tian, X. et al. Potent binding of 2019 novel coronavirus spike protein by a SARS coronavirus-specific human monoclonal antibody. Emerg Microbes Infect 9, 382–385 (2020).

6. Walls, A.C. et al. Structure, Function, and Antigenicity of the SARS-CoV-2 Spike Glycoprotein. Cell 181, 281-292.e6 (2020).

7. Liu, L. et al. Potent neutralizing antibodies against multiple epitopes on SARS-CoV-2 spike. Nature 584, 450–456 (2020).

8. Robbiani, D.F. et al. Convergent antibody responses to SARS-CoV-2 in convalescent individuals. Nature (2020).

9. Piccoli, L. et al. Mapping Neutralizing and Immunodominant Sites on the SARS-CoV-2 Spike Receptor-Binding Domain by Structure-Guided High-Resolution Serology. Cell 183, 1024–1042 e21 (2020).

10. Barnes, C.O. et al. Structures of human antibodies bound to SARS-CoV-2 spike reveal common epitopes and recurrent features of antibodies. bioRxiv (2020).

11. Greaney, A.J. et al. Complete Mapping of Mutations to the SARS-CoV-2 Spike Receptor-Binding Domain that Escape Antibody Recognition. Cell Host & Microbe 29, 44-57.e9 (2021).

12. Starr, T.N. et al. Deep Mutational Scanning of SARS-CoV-2 Receptor Binding Domain Reveals Constraints on Folding and ACE2 Binding. Cell 182, 1295–1310 e20 (2020).

13. Weisblum, Y. et al. Escape from neutralizing antibodies by SARS-CoV-2 spike protein variants. eLife 9, e61312 (2020).

14. Kemp, S.A. et al. SARS-CoV-2 evolution during treatment of chronic infection. Nature (2021).

15. Liu, Z. et al. Identification of SARS-CoV-2 spike mutations that attenuate monoclonal and serum antibody neutralization. Cell Host & Microbe (2021).

16. Willett, B.J. et al. The hyper-transmissible SARS-CoV-2 Omicron variant exhibits significant antigenic change, vaccine escape and a switch in cell entry mechanism. medRxiv, 2022.01.03.21268111 (2022).

17. Liu, L. et al. Striking antibody evasion manifested by the Omicron variant of SARS-CoV-2. Nature 602, 676–681 (2022).

18. Amanat, F. et al. SARS-CoV-2 mRNA vaccination induces functionally diverse antibodies to NTD, RBD, and S2. Cell 184, 3936–3948 e10 (2021).

19. Voss, W.N. et al. Prevalent, protective, and convergent IgG recognition of SARS-CoV-2 non-RBD spike epitopes in COVID-19 convalescent plasma. bioRxiv (2020).

20. Wang, Z. et al. Analysis of memory B cells identifies conserved neutralizing epitopes on the N-terminal domain of variant SARS-Cov-2 spike proteins. Immunity 55, 998–1012 e8 (2022).

21. Changrob, S. et al. Cross-Neutralization of Emerging SARS-CoV-2 Variants of Concern by Antibodies Targeting Distinct Epitopes on Spike. mBio 12, e0297521 (2021).

22. Chi, X. et al. A neutralizing human antibody binds to the N-terminal domain of the Spike protein of SARS-CoV-2. Science 369, 650–655 (2020).

23. London, S.D., Schmaljohn, A.L., Dalrymple, J.M. & Rice, C.M. Infectious enveloped RNA virus antigenic chimeras. Proc Natl Acad Sci U S A 89, 207–11 (1992).

24. Major, M.E. et al. DNA-based immunization with chimeric vectors for the induction of immune responses against the hepatitis C virus nucleocapsid. J Virol 69, 5798–805 (1995).

25. Muster, T. et al. Mucosal model of immunization against human immunodeficiency virus type 1 with a chimeric influenza virus. J Virol 69, 6678–86 (1995).

26. Kitson, J.D., Burke, K.L., Pullen, L.A., Belsham, G.J. & Almond, J.W. Chimeric polioviruses that include sequences derived from two independent antigenic sites of foot-and-mouth disease virus (FMDV) induce neutralizing antibodies against FMDV in guinea pigs. J Virol 65, 3068–75 (1991).

27. Li, S. et al. Chimeric influenza virus induces neutralizing antibodies and cytotoxic T cells against human immunodeficiency virus type 1. J Virol 67, 6659–66 (1993).

28. Nachbagauer, R. et al. A universal influenza virus vaccine candidate confers protection against pandemic H1N1 infection in preclinical ferret studies. NPJ Vaccines 2, 26 (2017).

29. Nachbagauer, R. et al. A chimeric hemagglutinin-based universal influenza virus vaccine approach induces broad and long-lasting immunity in a randomized, placebo-controlled phase I trial. Nat Med 27, 106–114 (2021).

30. Martinez, D.R. et al. Chimeric spike mRNA vaccines protect against Sarbecovirus challenge in mice. Science 373, 991–998 (2021).

31. Jumper, J. et al. Highly accurate protein structure prediction with AlphaFold. Nature 596, 583–589 (2021).

32. Jendrusch, M., Korbel, J.O. & Sadiq, S.K. AlphaDesign: A de novo protein design framework based on AlphaFold. bioRxiv, 2021.10.11.463937 (2021).

33. Moffat, L., Greener, J.G. & Jones, D.T. Using AlphaFold for Rapid and Accurate Fixed Backbone Protein Design. bioRxiv, 2021.08.24.457549 (2021).

34. Anishchenko, I. et al. De novo protein design by deep network hallucination. Nature 600, 547–552 (2021).

35. Lin, Z. et al. Evolutionary-scale prediction of atomic level protein structure with a language model. bioRxiv, 2022.07.20.500902 (2022).

36. Baek, M. et al. Accurate prediction of protein structures and interactions using a three-track neural network. Science 373, 871–876 (2021).

37. Abramson, J. et al. Accurate structure prediction of biomolecular interactions with AlphaFold 3. Nature 630, 493–500 (2024).

38. Schymkowitz, J. et al. The FoldX web server: an online force field. Nucleic Acids Res 33, W382–8 (2005).

39. Yin, S., Ding, F. & Dokholyan, N.V. Eris: an automated estimator of protein stability. Nat Methods 4, 466–7 (2007).

40. Rodrigues, C.H., Pires, D.E. & Ascher, D.B. DynaMut: predicting the impact of mutations on protein conformation, flexibility and stability. Nucleic Acids Res 46, W350–W355 (2018).

41. Quan, L., Lv, Q. & Zhang, Y. STRUM: structure-based prediction of protein stability changes upon single-point mutation. Bioinformatics 32, 2936–46 (2016).

42. Parthiban, V., Gromiha, M.M. & Schomburg, D. CUPSAT: prediction of protein stability upon point mutations. Nucleic Acids Res 34, W239–42 (2006).

43. Cheng, J., Randall, A. & Baldi, P. Prediction of protein stability changes for single-site mutations using support vector machines. Proteins 62, 1125–32 (2006).

44. Nuqui, X. et al. Simulation-driven design of stabilized SARS-CoV-2 spike S2 immunogens. Nature Communications 15, 7370 (2024).

45. Mariani, V., Biasini, M., Barbato, A. & Schwede, T. lDDT: a local superposition-free score for comparing protein structures and models using distance difference tests. Bioinformatics 29, 2722–8 (2013).

46. Hsieh, C.L. et al. Structure-based Design of Prefusion-stabilized SARS-CoV-2 Spikes. Science 369, 1501–1505 (2020).

47. Rice, P., Longden, I. & Bleasby, A. EMBOSS: the European Molecular Biology Open Software Suite. Trends Genet 16, 276–7 (2000).

48. Cai, Y. et al. Distinct conformational states of SARS-CoV-2 spike protein. Science 369, 1586 (2020).

49. Abramson, J. et al. Accurate structure prediction of biomolecular interactions with AlphaFold 3. Nature (2024).

50. Wrapp, D. et al. Cryo-EM structure of the 2019-nCoV spike in the prefusion conformation. Science 367, 1260–1263 (2020).

51. Huang, Y., Yang, C., Xu, X.F., Xu, W. & Liu, S.W. Structural and functional properties of SARS-CoV-2 spike protein: potential antivirus drug development for COVID-19. Acta Pharmacol Sin 41, 1141–1149 (2020).

52. Edgar, R.C. MUSCLE: multiple sequence alignment with high accuracy and high throughput. Nucleic Acids Res 32, 1792–7 (2004).

53. Evans, R. et al. Protein complex prediction with AlphaFold-Multimer. bioRxiv, 2021.10.04.463034 (2022).

54. Pronk, S. et al. GROMACS 4.5: a high-throughput and highly parallel open source molecular simulation toolkit. Bioinformatics 29, 845–54 (2013).

55. Huang, J. & MacKerell, A.D., Jr. CHARMM36 all-atom additive protein force field: validation based on comparison to NMR data. J Comput Chem 34, 2135–45 (2013).

56. Bussi, G., Donadio, D. & Parrinello, M. Canonical sampling through velocity rescaling. J Chem Phys 126, 014101 (2007).

57. Sengar, A., Cervantes, M., Bondalapati, S.T., Hess, T. & Kasson, P.M. Single-virus fusion measurements reveal multiple mechanistically equivalent pathways for SARS-CoV-2 entry. Journal of Virology, e01992–22 (2023).

58. Crawford, K.H.D. et al. Protocol and Reagents for Pseudotyping Lentiviral Particles with SARS-CoV-2 Spike Protein for Neutralization Assays. Viruses 12(2020).

