## Supplementary Information for "Structural prediction of chimeric immunogens to elicit targeted antibodies against betacoronaviruses"

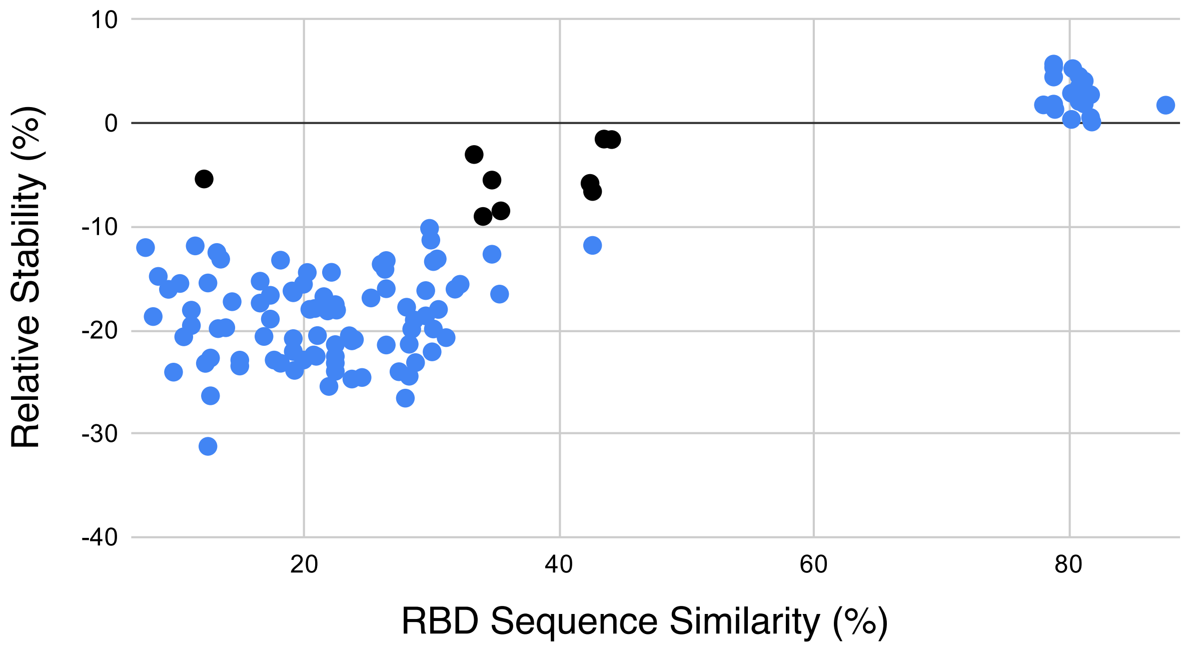


**Figure S1. Predicted Stability of Coronavirus RBD Chimeras**. The relative stability score for each receptor-binding domain (RBD) chimera is plotted against the sequence similarity between the parental RBD sequence and SARS-CoV-2. There is less of a clear low-similarity, high-stability cluster than for the S1/S2 chimeras plotted in Fig. 3. However, the 9 sequences with similarity <45% and relative stability >-10% were selected for further analysis by molecular dynamics simulation.

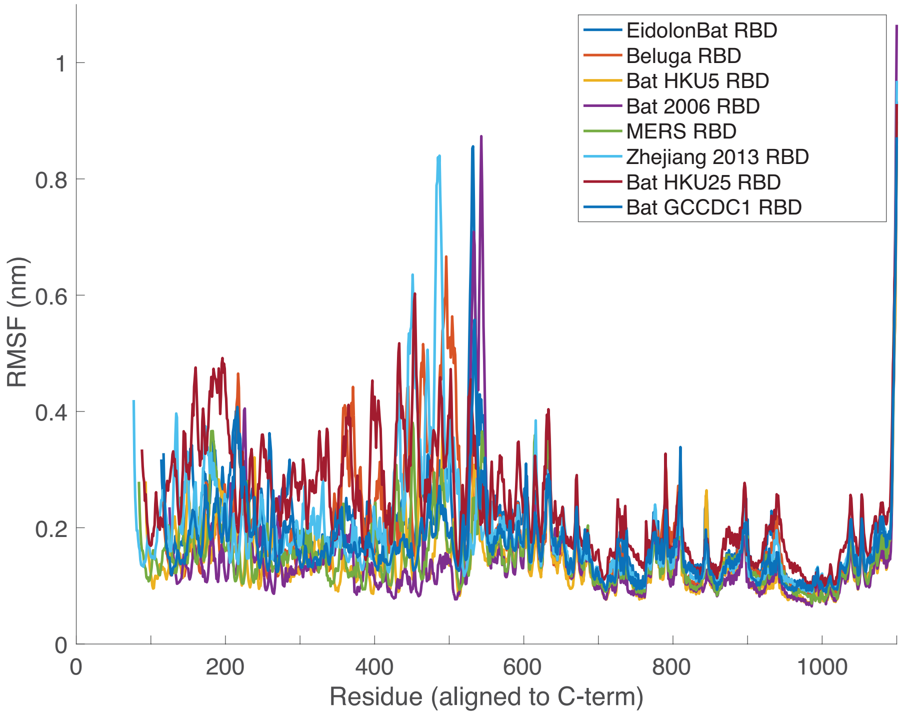


**Figure S2. Per-residue structural stability of simulated RBD chimeras.** Root-mean-squared fluctuation (RMSF) values are plotted per residue for each of the simulated chimeras. Values were calculated on nanosecond intervals throughout the simulation trajectory.

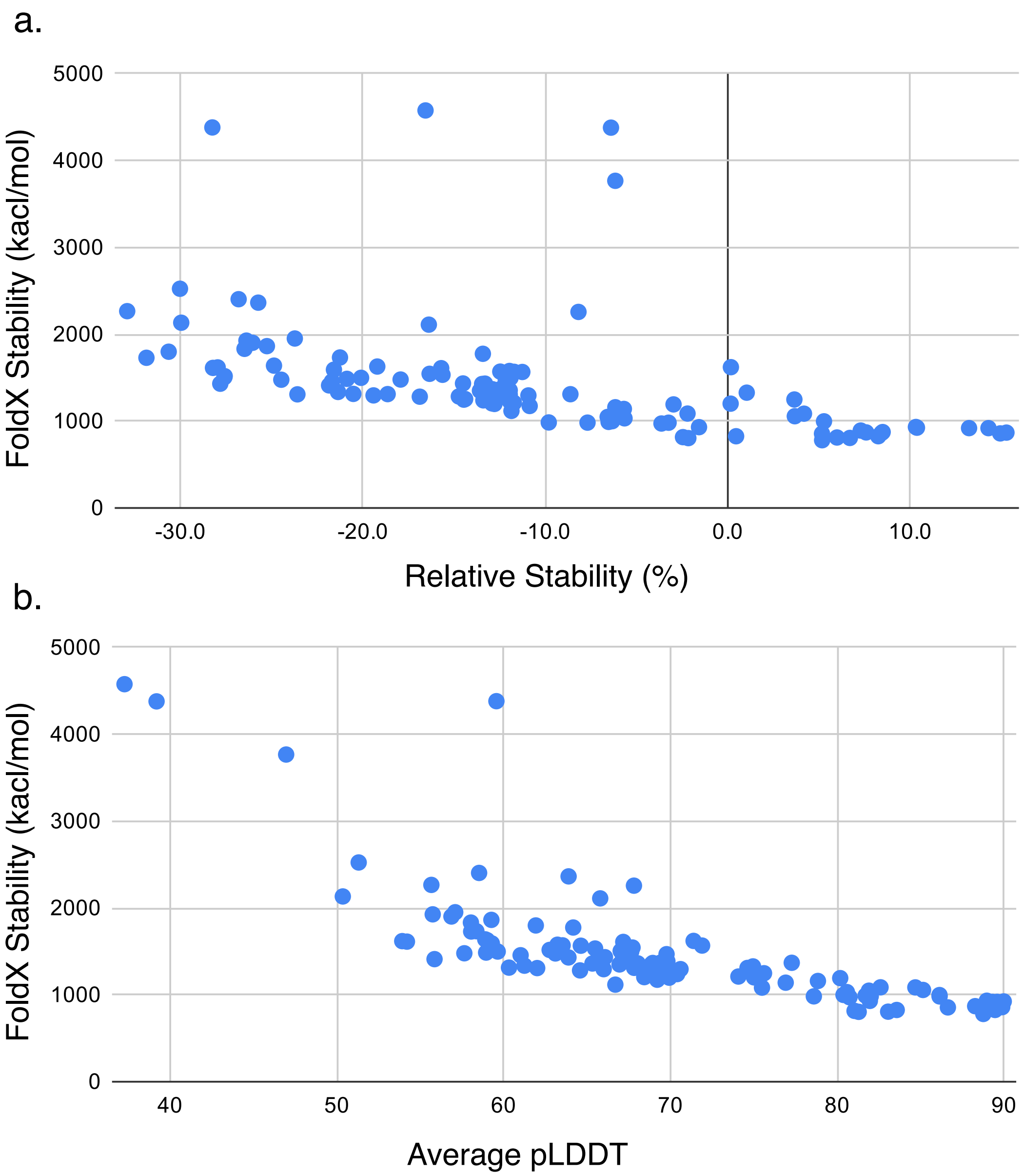


**Figure S3. Comparison of FoldX Scores with AlphaFold pLDDT and Relative Stability Scores.** FoldX stability scores were calculated for each chimera and compared both to the relative stability metric (a) and the average pLDDT output from AlphaFold (b). Overall trends were similar, with a Spearman rank correlation of -0.89 between FoldX stability score and AlphaFold pLDDT and -0.78 between FoldX stability score and relative stability. Both plots show outliers with substantially higher FoldX stability.

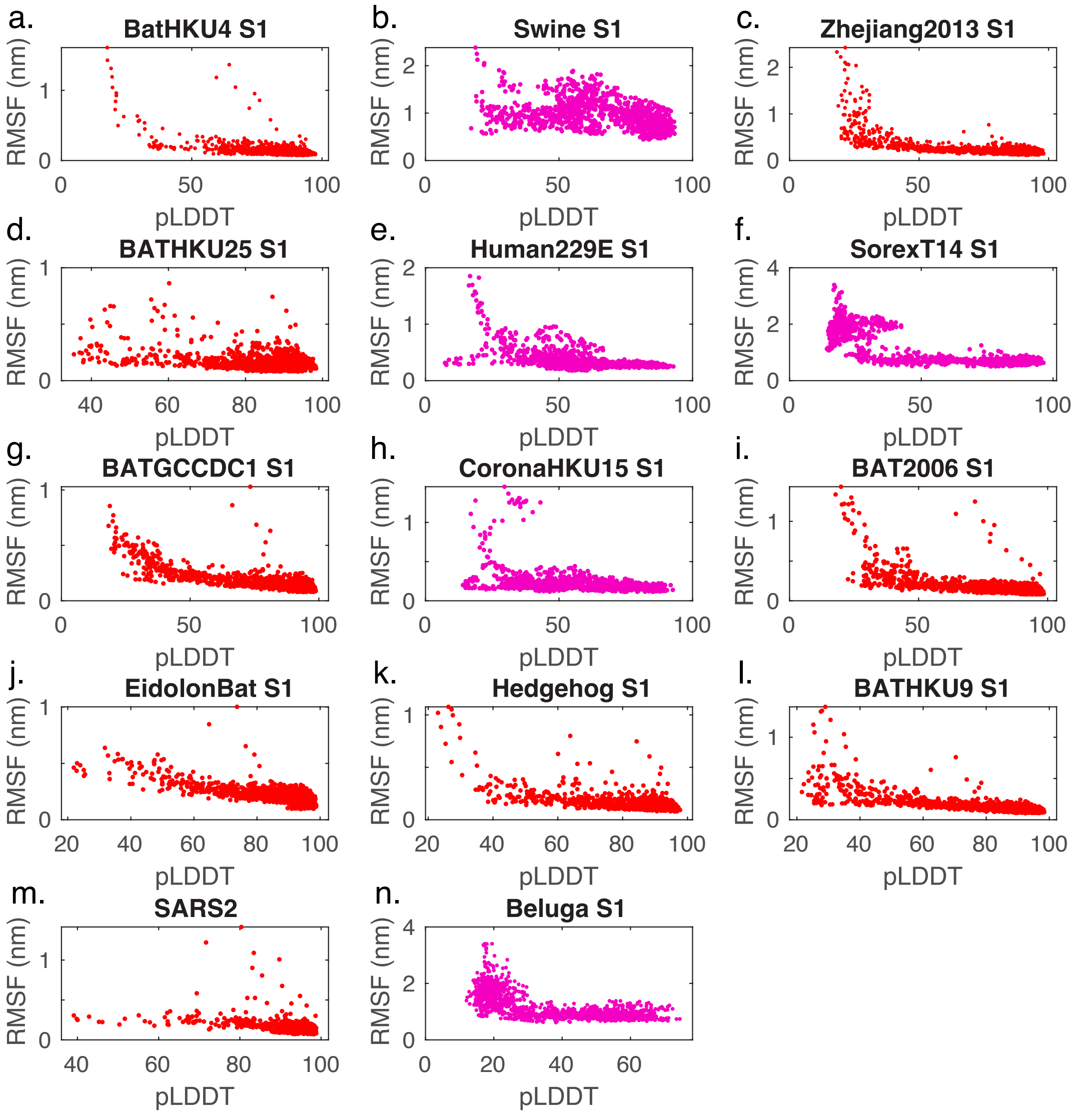


**Figure S4. Plots of pLDDT versus RMSF over all simulated S1 chimeras**. Panels (a-n) show the per-residue pLDDT from AlphaFold plotted against the RMSF from molecular dynamics simulations across the entire dataset of simulated S1 chimeras. Predicted high-stability chimeras are plotted in red, and predicted low-stability chimeras are plotted in magenta. The Spearman correlation coefficient between pLDDT and RMSF over the dataset is -0.69.

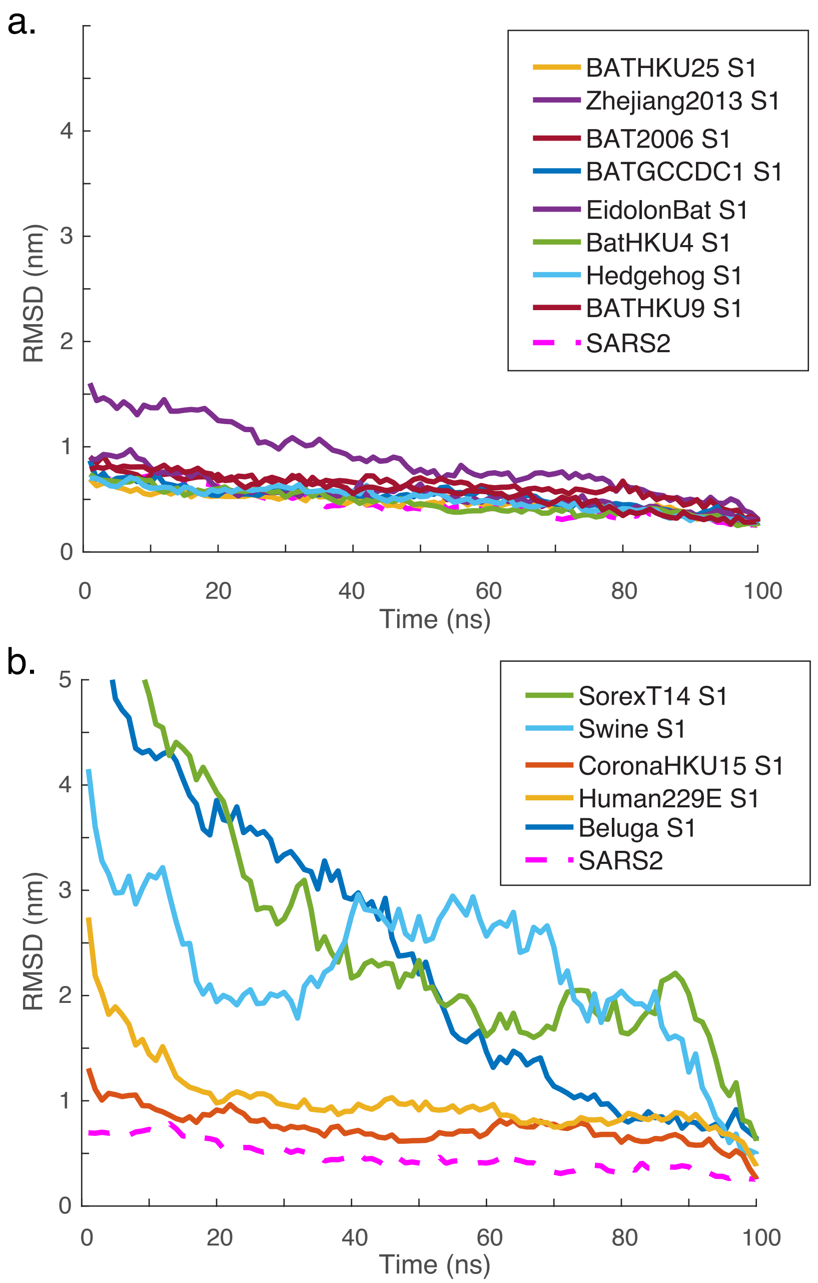


**Figure S5. Structural stability of predicted chimeras relative to the end of the simulation.** Root-mean-squared deviation values are plotted relative to simulated structures after 100 ns of simulation. Panel (a) shows the predicted-stable chimeras, and panel (b) shows the predicted-unstable chimeras.

| Accession ID | Full Name | Short Name |
|---|---|---|
| QWN56273 | Alphacoronavirus sp. | Alpha |
| YP_009199242 | Alphacoronavirus 1 | Alpha1 |
| QDF43810 | Coronavirus BtRs-AlphaCoV/YN2018 | AlphaCorona |
| YP_009200735 | BtRf-AlphaCoV/YN2012 | AlphaCorona2012 |
| AIA62242 | BtMf-AlphaCoV/HuB2013-a | AlphaCorona2013 |
| AIA62240 | BtMf-AlphaCoV/GD2012-a | AlphaCoronaGD2012 |
| QXT50810 | Anser fabalis coronavirus NCN2 | Anser |
| UBB42470 | Jingmen Apodemus agrarius alphacoronavirus 1 | ApodemusAlpha |
| UBB42425 | Jingmen Apodemus agrarius betacoronavirus 1 | ApodemusBeta |
| YP_009824998 | Avian coronavirus | Avian |
| ADX59474 | Rousettus bat coronavirus/Kenya/KY06/2006 | BAT2006 |
| ALJ94036 | SARS-like coronavirus BatCoV/BB9904/BGR/2008 | BAT2008 |
| QHA24665 | Rousettus aegyptiacus bat coronavirus 229E-related | BAT229E |
| ACU31051 | Bat SARS Cov Rs806/2006 | Bat806 |
| ABG11963 | Bat coronavirus A515/2005 | BatA515 |
| ABG11964 | Bat coronavirus A527/2005 | BatA527 |
| ABG11965 | Bat coronavirus A701/2005 | BatA701 |
| QLE11825 | Bat alphacoronavirus | BatAlpha |
| YP_003858584 | Bat coronavirus BM48-31/BGR/2008 | BatBGR |
| QHA24724 | Hipposideros pomona bat coronavirus CHB25 | BATCHB25 |
| YP_009273005 | Rousettus bat coronavirus GCCDC1 | BATGCCDC1 |
| YP_008439202 | Bat coronavirus CDPHE15 | BatHE15 |
| QHA24710 | Hipposideros pomona bat coronavirus HKU10-related | BATHKU10rel |
| YP_006908642 | Bat coronavirus HKU10 | BatHKU10 |
| ASL68941 | Hypsugo bat coronavirus HKU25 | BATHKU25 |
| QCX35160 | Tylonycteris bat coronavirus HKU33 | BatHKU33 |
| YP_001039953 | Tylonycteris bat coronavirus HKU4 | BatHKU4 |
| YP_001039962 | Pipistrellus bat coronavirus HKU5 | BATHKU5 |
| YP_001039971 | Rousettus bat coronavirus HKU9 | BATHKU9 |
| ADX59495 | Chaerephon bat coronavirus/Kenya/KY22/2006 | BatKY22 |
| ADX59458 | Chaerephon bat coronavirus/Kenya/KY41/2006 | BatKY41 |
| ADX59451 | Cardioderma bat coronavirus/Kenya/KY43/2006 | BatKY43 |
| YP_001876437 | Beluga whale coronavirus SW1 | Beluga |
| AYR18599 | Betacoronavirus sp. | Beta |
| QWN56232 | Betacoronavirus sp. RsYN03 | Beta03 |
| QWN56242 | Betacoronavirus sp. RsYN04 | Beta04 |
| QWN56202 | Betacoronavirus sp. RmYN05 | Beta05 |
| QWN56252 | Betacoronavirus sp. RpYN06 | Beta06 |

| | | |
|---|---|---|
| QWN56212 | Betacoronavirus sp. RmYN07 | Beta07 |
| QWN56222 | Betacoronavirus sp. RmYN08 | Beta08 |
| QWN56263 | Betacoronavirus sp. RsYN09 | Beta09 |
| YP_009555241 | Betacoronavirus 1 | Beta1 |
| AGC51116 | Betacoronavirus BtCoV/KW2E-F93/Nyc_spec/GHA/2010 | Beta2010 |
| QJX58373 | Coronavirus BtRt-BetaCoV/GX2018 | BetaCorona |
| AIA62340 | BtRf-BetaCoV/HuB2013 | BetaCorona2013 |
| QDF43815 | Coronavirus BtRl-BetaCoV/SC2018 | BetaCoronaSC2018 |
| QRN68024 | Betacoronavirus Erinaceus | BetaErin |
| YP_002308479 | Bulbul coronavirus HKU11 | Bulbul |
| ATI09449 | Camel coronavirus HKU23 | Camel |
| YP_009513021 | Coronavirus HKU15 | CoronaHKU15 |
| YP_009380521 | Coronavirus AcCoV-JC34 | CoronaJC34 |
| AWR88316 | Deltacoronavirus sp. | Delta |
| QII89019 | Bottlenose dolphin coronavirus | Dolphin |
| ADX59466 | Eidolon bat coronavirus/Kenya/KY24/2006 | EidolonBat |
| BBC54822 | Falcon coronavirus UAE-HKU27 | FalconHKU27 |
| AYF53093 | Feline alphacoronavirus 1 | FelineAlpha |
| YP_009256197 | Ferret coronavirus | Ferret |
| ABO88150 | Bat coronavirus Fujian/773/2005 | Fujian |
| YP_009755897 | Canada goose coronavirus | Goose |
| YP_009513010 | Hedgehog coronavirus 1 | Hedgehog |
| QGA70692 | Erinaceus hedgehog coronavirus HKU31 | HedgehogHKU27 |
| YP_005352863 | Night heron coronavirus HKU19 | HeronHKU19 |
| BBC54832 | Houbara coronavirus UAE-HKU28 | HoubaraHKU28 |
| YP_009194639 | Human coronavirus 229E | Human229E |
| YP_173238 | Human coronavirus HKU1 | HumanHKU1 |
| YP_003767 | Human coronavirus NL63 | HumanNL63 |
| AZF86130 | Alphacoronavirus Bat-CoV/P.kuhlii/Italy/206679-3/2010 | Italy2010 |
| AZF86118 | Alphacoronavirus Bat-CoV/P.kuhlii/Italy/206645-41/2011 | Italy2011 |
| YP_009755890 | Alphacoronavirus Bat-CoV/P.kuhlii/Italy/3398-19/2015 | Italy2015 |
| UBB42440 | Jingmen Myotis chinensis alphacoronavirus 1 | JingmenAlpha |
| QVN46559 | Bat SARS-like coronavirus Khosta-1 | Khosta1 |
| QVN46569 | Bat SARS-like coronavirus Khosta-2 | Khosta2 |
| UBB42462 | Longquan Berylmys bowersi alphacoronavirus 1 | LongquanAlpha |
| UBB42416 | Longquan Niviventer niviventer betacoronavirus 1 | LongquanBeta |
| QOE77336 | Longquan Rl rat coronavirus | LongquanRat |
| YP_009336484 | Lucheng Rn rat coronavirus | LuchengRat |
| YP_005352854 | Magpie-robin coronavirus HKU18 | Magpie |
| YP_009047204 | Middle East respiratory syndrome-related coronavirus | MERS |
| QPP46979 | Minacovirus mink/NLD/2020/NT_4 | Minacovirus2020 |
| YP_001718605 | Miniopterus bat coronavirus 1 | Minio |

| | | |
|---|---|---|
| ADX59482 | Miniopterus bat coronavirus/Kenya/KY27/2006 | Minio2006 |
| UBB42447 | Jingmen Miniopterus schreibersii alphacoronavirus 1 | MinioAlpha |
| YP_001718612 | Miniopterus bat coronavirus HKU8 | MinioHKU8 |
| QHA24671 | Miniopterus pusillus bat coronavirus HKU8-related | MinioHKU8rel |
| YP_009019182 | Mink coronavirus 1 | Mink |
| AVY53336 | Alphacoronavirus Mink/China/1/2016 | Mink2016 |
| ADI80523 | Mink coronavirus strain WD1133 | MinkWD1133 |
| AFD29244 | Common moorhen coronavirus HKU21 | MoorHKU21 |
| YP_002308506 | Munia coronavirus HKU13 | Munia |
| YP_009824982 | Murine coronavirus | Murine |
| YP_009199609 | Myotis ricketti alphacoronavirus Sax-2011 | Myotis |
| AHN92552 | Mystacina coronavirus New Zealand/2013 | Mystacina |
| YP_009201730 | Nyctalus velutinus alphacoronavirus SC-2013 | NyctalusAlpha |
| QLR06864 | Pangolin coronavirus | Pangolin |
| QPN00063 | Apodemus peninsulae coronavirus | Peninsuale |
| BBC54842 | Pigeon coronavirus UAE-HKU29 | PigeonHKU29 |
| NP_598310 | Porcine epidemic diarrhea virus | PorcineVirus |
| AXP20281 | Quail deltacoronavirus | QuailDelta |
| BBC54852 | Quail coronavirus UAE-HKU30 | QuailHKU30 |
| YP_005454245 | Rabbit coronavirus HKU14 | RabbitHKU14 |
| YP_009113025 | China Rattus coronavirus HKU24 | RattusHKU24 |
| YP_009199790 | Rhinolophus ferrumequinum alphacoronavirus HuB-2013 | RhinoAlpha |
| UBB42405 | Jingmen Rhinolophus sinicus betacoronavirus 1 | RhinoBeta |
| YP_001552236 | Rhinolophus bat coronavirus HKU2 | RhinoHKU2 |
| QCX35167 | Rhinolophus bat coronavirus HKU32 | RhinoHKUU32 |
| UAL80443 | Swine acute diarrhea syndrome coronavirus | SADS |
| QZX47235 | Sarbecovirus sp. | Sarbecovirus |
| QYC92806 | Sarbecovirus RhGB01 | SarbecovirusRhG |
| YP_001351684 | Scotophilus bat coronavirus 512 | Scotophilus512 |
| ABO88151 | Bat coronavirus Shandong/977/2006 | Shandong |
| QPB10668 | Shorebird deltacoronavirus | ShorebirdDelta |
| ATP66784 | Sorex araneus coronavirus T14 | SorexT14 |
| AWV67107 | Sparrow deltacoronavirus | SparrowDelta |
| YP_005352846 | Sparrow coronavirus HKU17 | SparrowHKU17 |
| QEH62669 | Swine enteric alphacoronavirus | Swine |
| QZQ78890 | Tapir coronavirus 1044512-1 | Tapir |
| YP_002308497 | Thrush coronavirus HKU12-600 | ThrushHKU12 |
| QBG64657 | Alphacoronavirus UKRn3 | UKRn3 |
| YP_009824974 | Wencheng Sm shrew coronavirus | Wencheng |
| UBB42478 | Wenzhou Suncus murinus alphacoronavirus 1 | WenzhouAlpha |
| UBB42431 | Wenzhou Pipistrellus abramus betacoronavirus 1 | WenzhouBeta |
| YP_005352838 | White-eye coronavirus HKU16 | WhiteHKU16 |

| | | |
|---|---|---|
| YP_005352871 | Wigeon coronavirus HKU20 | WigeonHKU20 |
| ALK02457 | SARS-like coronavirus WIV16 | WIV16 |
| YP_009072440 | Bat Hp-betacoronavirus Zhejiang2013 | Zhejiang2013 |

**Table S1. Sequences used in this study.**

| Accession ID | Full Name | Short Name |
|---|---|---|
| AIA62240 | BtMf-AlphaCoV/GD2012-a | AlphaCoronaGD2012 |
| UBB42425 | Jingmen Apodemus agrarius betacoronavirus 1 | ApodemusBeta |
| ABG11963 | Bat coronavirus A515/2005 | BatA515 |
| ABG11964 | Bat coronavirus A527/2005 | BatA527 |
| ABG11965 | Bat coronavirus A701/2005 | BatA701 |
| QCX35160 | Tylonycteris bat coronavirus HKU33 | BatHKU33 |
| AGC51116 | Betacoronavirus BtCoV/KW2E-F93/Nyc_spec/GHA/2010 | Beta2010 |
| QJX58373 | Coronavirus BtRt-BetaCoV/GX2018 | BetaCorona |
| AIA62340 | BtRf-BetaCoV/HuB2013 | BetaCorona2013 |
| ATI09449 | Camel coronavirus HKU23 | Camel |
| UBB42478 | Wenzhou Suncus murinus alphacoronavirus 1 | WenzhouAlpha |
| UBB42431 | Wenzhou Pipistrellus abramus betacoronavirus 1 | WenzhouBeta |

**Table S2: Sequences Excluded due to AlphaFold Failures**

| Source for RBD (short name) | RBD Sequence Similarity (%) | Relative Stability (%) |
|---|---|---|
| Alpha | 20.7 | -22.4 |
| Alpha1 | 12.6 | -26.3 |
| AlphaCorona | 20.2 | -14.4 |
| AlphaCorona2012 | 30.5 | -18.0 |
| AlphaCorona2013 | 12.6 | -22.7 |
| Anser | 26.3 | -14.1 |
| ApodemusAlpha | 26 | -13.6 |
| Avian | 20.4 | -18.0 |
| BAT2006 | 42.6 | -6.6 |
| BAT2008 | 80.2 | 0.4 |
| BAT229E | 22.4 | -23.2 |
| Bat806 | 81.2 | 4.1 |
| BatAlpha | 8.1 | -18.7 |
| BatBGR | 80.2 | 2.9 |
| BATCHB25 | 30.1 | -19.9 |
| BATGCCDC1 | 42.4 | -5.8 |
| BatHE15 | 17.6 | -22.9 |

| | | |
|---|---|---|
| BatHKU10 | 19.1 | -16.4 |
| BATHKU10 | 14.3 | -17.2 |
| BATHKU25 | 33.3 | -3.0 |
| BatHKU33 | 19 | -16.2 |
| BatHKU4 | 34.7 | -12.7 |
| BATHKU5 | 34.7 | -5.5 |
| BATHKU9 | 42.6 | -11.8 |
| BatKY22 | 31.8 | -16.0 |
| BatKY41 | 22.4 | -17.5 |
| BatKY43 | 27.9 | -26.6 |
| Beluga | 12.1 | -5.4 |
| Beta | 23.5 | -20.5 |
| Beta03 | 80.8 | 4.5 |
| Beta04 | 78.8 | 4.5 |
| Beta05 | 78.8 | 5.4 |
| Beta06 | 81.2 | 1.9 |
| Beta07 | 80.8 | 3.5 |
| Beta08 | 78.8 | 5.7 |
| Beta09 | 78.8 | 1.8 |
| Beta1 | 24.5 | -24.6 |
| BetaCoronaSC2018 | 81.7 | 0.5 |
| BetaErin | 29.8 | -10.2 |
| Bulbul | 16.8 | -20.6 |
| CoronaHKU15 | 16.5 | -15.3 |
| CoronaJC34 | 26.4 | -13.3 |
| Delta | 13.4 | -13.1 |
| Dolphin | 22.5 | -18.1 |
| EidolonBat | 44.1 | -1.6 |
| FalconHKU27 | 8.5 | -14.8 |
| FelineAlpha | 14.9 | -23.5 |
| Ferret | 19.1 | -22.1 |
| Fujian | 21.9 | -25.4 |
| Goose | 21.8 | -18.1 |
| Hedgehog | 29.9 | -11.3 |
| HedgehogHKU27 | 34 | -9.0 |
| HeronHKU19 | 32.2 | -15.6 |
| HoubaraHKU28 | 22.4 | -21.4 |
| Human229E | 29.5 | -16.2 |
| HumanHKU1 | 26.4 | -16.0 |
| HumanNL63 | 22.4 | -22.5 |
| Italy2010 | 16.5 | -17.4 |
| Italy2011 | 28.2 | -21.4 |

| | | |
|---|---|---|
| Italy2015 | 13.8 | -19.8 |
| JingmenAlpha | 17.3 | -16.6 |
| Khosta1 | 80.3 | 5.3 |
| Khosta2 | 78 | 1.8 |
| LongquanAlpha | 31.1 | -20.7 |
| LongquanBeta | 26.4 | -21.4 |
| LongquanRat | 25.2 | -16.9 |
| LuchengRat | 28.4 | -19.9 |
| Magpie | 13.1 | -12.5 |
| MERS | 35.4 | -8.5 |
| Minacovirus2020 | 11.1 | -19.5 |
| Minio | 13.2 | -19.8 |
| Minio2006 | 14.9 | -22.9 |
| MinioAlpha | 21 | -20.5 |
| MinioHKU8 | 10.2 | -15.5 |
| MinioHKU8rel | 18.1 | -23.2 |
| Mink | 11.1 | -18.1 |
| Mink2016 | 20.8 | -17.9 |
| MinkWD1133 | 12.2 | -23.2 |
| MoorHKU21 | 29.5 | -18.6 |
| Munia | 21.5 | -16.8 |
| Murine | 28.6 | -19.0 |
| Myotis | 17.3 | -18.9 |
| Mystacina | 9.7 | -24.1 |
| NyctalusAlpha | 20.9 | -22.5 |
| Pangolin | 87.6 | 1.7 |
| Peninsuale | 28 | -17.8 |
| PigeonHKU29 | 22.4 | -24.0 |
| PorcineVirus | 12.4 | -31.2 |
| QuailDelta | 12.4 | -15.4 |
| QuailHKU30 | 7.5 | -12.0 |
| RabbitHKU14 | 23.7 | -24.7 |
| RattusHKU24 | 28.7 | -23.1 |
| RhinoAlpha | 19.9 | -22.9 |
| RhinoBeta | 80.8 | 2.1 |
| RhinoHKU2 | 30 | -22.1 |
| RhinoHKUU32 | 10.5 | -20.6 |
| SADS | 28.2 | -24.4 |
| Sarbecovirus | 81.7 | 2.8 |
| SarbecovirusRhG | 78.9 | 1.3 |
| Scotophilus512 | 19.2 | -23.9 |
| Shandong | 19.1 | -20.8 |

| | | |
|---|---|---|
| ShorebirdDelta | 9.3 | -16.1 |
| SorexT14 | 22.1 | -14.4 |
| SparrowDelta | 18.1 | -13.2 |
| SparrowHKU17 | 19.9 | -15.6 |
| Swine | 27.4 | -24.0 |
| Tapir | 23.7 | -21.0 |
| ThrushHKU12 | 35.3 | -16.5 |
| UKRn3 | 30.1 | -13.4 |
| Wencheng | 23.9 | -20.9 |
| WhiteHKU16 | 11.4 | -11.9 |
| WigeonHKU20 | 30.4 | -13.1 |
| WIV16 | 81.8 | 0.1 |
| Zhejiang2013 | 43.5 | -1.5 |

**Table S3: Relative Stability Scores and Sequence Similarity for RBD Chimeras**. Names for chimeras selected for MD simulation are colored in red.

| Source for S1 (short name) | S1 Sequence Similarity (%) | Relative Stability (%) |
|---|---|---|
| RhinoBeta | 71.8 | 15.3 |
| Beta03 | 72.2 | 14.9 |
| Bat806 | 72.8 | 14.3 |
| Beta07 | 71.8 | 13.2 |
| Pangolin | 78.6 | 10.4 |
| Beta06 | 74.6 | 10.3 |
| SarbecovirusRhG | 68.5 | 8.5 |
| BatBGR | 69.1 | 8.4 |
| Khosta2 | 70.6 | 8.2 |
| BAT2008 | 67.8 | 7.6 |
| Sarbecovirus | 72.6 | 7.3 |
| EidolonBat | 35.9 | 6.7 |
| Khosta1 | 69.8 | 6.0 |
| Beta09 | 70.1 | 5.3 |
| BetaCoronaSC2018 | 73.2 | 5.2 |
| WIV16 | 72 | 5.2 |
| Beta08 | 64.6 | 4.2 |
| Beta05 | 64.6 | 3.7 |
| BATGCCDC1 | 35.9 | 3.7 |
| BAT2006 | 36.6 | 1.0 |
| BATHKU25 | 32.1 | 0.5 |
| Zhejiang2013 | 39.3 | 0.2 |
| BATHKU9 | 38.1 | 0.2 |

| | | |
|---|---|---|
| BatHKU4 | 32.2 | -1.6 |
| Hedgehog | 35.5 | -2.2 |
| Beta04 | 64.6 | -2.2 |
| BetaErin | 35.1 | -2.5 |
| HumanHKU1 | 32.8 | -3.0 |
| MERS | 32.4 | -3.2 |
| HedgehogHKU27 | 34.9 | -3.6 |
| Tapir | 33.2 | -5.7 |
| Peninsuale | 35.6 | -5.7 |
| Murine | 35.4 | -6.2 |
| SorexT14 | 30.8 | -6.2 |
| LongquanBeta | 33.4 | -6.2 |
| BATHKU5 | 33.2 | -6.3 |
| Dolphin | 21.5 | -6.4 |
| LongquanRat | 33.3 | -6.5 |
| RabbitHKU14 | 33.7 | -6.6 |
| Beta | 35.3 | -7.7 |
| FelineAlpha | 21 | -8.2 |
| Alpha1 | 24 | -8.6 |
| Beta1 | 32.2 | -9.8 |
| RhinoHKUU32 | 26.6 | -10.9 |
| Ferret | 21.3 | -10.9 |
| BatAlpha | 24.6 | -11.3 |
| PorcineVirus | 24.4 | -11.7 |
| RattusHKU24 | 34.9 | -11.7 |
| MinkWD1133 | 24.6 | -11.9 |
| Italy2010 | 25.4 | -11.9 |
| RhinoAlpha | 23.2 | -12.0 |
| MinioHKU8 | 25.8 | -12.0 |
| BatKY43 | 24.6 | -12.0 |
| Minacovirus2020 | 22.4 | -12.0 |
| Mink2016 | 22.7 | -12.1 |
| BATHKU10 | 25.2 | -12.1 |
| MinioAlpha | 25.4 | -12.2 |
| HumanNL63 | 25.3 | -12.3 |
| Italy2011 | 23.8 | -12.4 |
| AlphaCorona2013 | 24.1 | -12.5 |
| Mink | 22.9 | -12.5 |
| BatHKU33 | 25.2 | -12.8 |
| BATCHB25 | 25.4 | -12.9 |
| BatKY22 | 22.4 | -12.9 |
| BatHE15 | 25.6 | -13.3 |

| | | |
|---|---|---|
| MinioHKU8rel | 20.5 | -13.3 |
| Shandong | 21.4 | -13.4 |
| Scotophilus512 | 25.6 | -13.4 |
| Fujian | 24.2 | -13.4 |
| BatKY41 | 23.4 | -13.5 |
| Italy2015 | 25.2 | -13.6 |
| Minio | 25.7 | -14.4 |
| NyctalusAlpha | 23 | -14.5 |
| Alpha | 25.5 | -14.5 |
| Myotis | 26.7 | -14.7 |
| AlphaCorona | 22.7 | -15.6 |
| JingmenAlpha | 27.1 | -15.7 |
| Minio2006 | 19.8 | -16.3 |
| BatHKU10 | 26.8 | -16.4 |
| Beluga | 23 | -16.6 |
| ShorebirdDelta | 30.8 | -16.9 |
| Delta | 32 | -17.9 |
| Mystacina | 24.7 | -18.6 |
| ThrushHKU12 | 32.9 | -19.2 |
| BAT229E | 21.7 | -19.4 |
| QuailDelta | 30.9 | -20.1 |
| CoronaJC34 | 32.5 | -20.5 |
| LuchengRat | 25.1 | -20.9 |
| UKRn3 | 30.7 | -21.3 |
| ApodemusAlpha | 32.9 | -21.4 |
| WhiteHKU16 | 31.4 | -21.6 |
| Anser | 29.6 | -21.7 |
| Wencheng | 29 | -21.9 |
| Bulbul | 27.6 | -23.6 |
| WigeonHKU20 | 29.9 | -23.7 |
| Munia | 29.7 | -24.5 |
| Magpie | 28 | -24.9 |
| PigeonHKU29 | 30.2 | -25.3 |
| Avian | 27.5 | -25.7 |
| QuailHKU30 | 32 | -26.1 |
| SparrowHKU17 | 23.7 | -26.4 |
| HoubaraHKU28 | 29.9 | -26.5 |
| FalconHKU27 | 31.1 | -26.8 |
| SADS | 30.3 | -27.6 |
| RhinoHKU2 | 33.7 | -27.6 |
| AlphaCorona2012 | 29.9 | -27.8 |
| Goose | 29 | -28.0 |

| | | |
|---|---|---|
| LongquanAlpha | 26 | -28.2 |
| SparrowDelta | 31.1 | -28.3 |
| HeronHKU19 | 28.3 | -30.0 |
| MoorHKU21 | 28.7 | -30.0 |
| Swine | 33.7 | -30.7 |
| Human229E | 31.7 | -31.9 |
| CoronaHKU15 | 31.7 | -32.9 |

**Table S4: Relative Stability Scores and Sequence Similarity for S1 Chimeras**. Names for predicted high-stability chimeras selected for MD simulation are colored in red, and names for predicted low-stability chimeras selected for MD simulation are colored in magenta.
